# cBAF drives AR-V7 expression through regulation of transcriptional termination in castration-resistant prostate cancer

**DOI:** 10.64898/2026.10.13.758074

**Authors:** Masataka Nakano, Shakhzodjon Uzokboev, Jacob G. Kirkland, W. Andy Tao, Emily C. Dykhuizen

## Abstract

The androgen receptor splice variant AR-V7 drives resistance to androgen receptor-targeted therapies in castration-resistant prostate cancer (CRPC), but the mechanisms governing its expression remain incompletely understood. Here, we identify the canonical BAF (cBAF) chromatin remodeling complex as a selective regulator of AR-V7 expression. Pharmacologic and genetic disruption of cBAF reduced AR-V7 levels, suppressed AR-V7-dependent transcriptional programs, and impaired CRPC cell proliferation. Mechanistically, cBAF binds an intronic regulatory element downstream of AR cryptic exon 3, where it maintains chromatin accessibility, enhancer activity, and promoter-enhancer looping. cBAF loss decreases recruitment of transcription termination machinery, reduces RNA polymerase II Thr4 phosphorylation, and promotes transcriptional readthrough beyond the AR-V7 polyadenylation site. POINT-seq revealed that cBAF-dependent transcriptional termination is required for AR-V7 biogenesis and regulates termination at a subset of genes genome-wide. Together, these findings uncover a role for cBAF in chromatin-mediated control of transcriptional termination and establish this mechanism as a critical determinant of AR-V7 expression.

**Key Findings:**

- cBAF is the only SWI/SNF subcomplex required for AR-V7 expression and function.
- AR-V7 transcription requires cBAF-promoted premature transcriptional termination in addition to alternative splicing.
- cBAF maintains an intronic enhancer-like regulatory element through chromatin accessibility, H3K27ac, and chromatin looping.
- cBAF broadly influences RNA polymerase II termination dynamics, revealing a previously unrecognized role in transcriptional termination.

## Introduction

Prostate cancer (PC) remains a leading cause of cancer-related morbidity and mortality in men worldwide, with steadily increasing incidence (Siegel et al., 2023). PC is largely driven by androgen receptor (AR)-dependent transcriptional programs (Wong et al., 2014), making androgen deprivation and AR antagonists the standard first-line treatments (Rebello et al., 2014; Sternberg et al., 2020). While these treatments are initially very effective, most PCs eventually progress to castration-resistant prostate cancer (CRPC), in which AR signaling is reactivated despite androgen depletion (Scher and Sawyers, 2005). This reactivation is mediated through multiple mechanisms, including AR gene amplification, AR-activating mutations, increased intratumoral androgen synthesis, and overexpression of AR co-regulators (Linja et al., 2001; Robinson et al., 2015; Cato et al., 2017). A substantial fraction (∼20-30% (Del Re et al., 2019)) of CRPC expresses AR-V7, a transcript variant of AR (Kanayama et al., 2021). Canonical full-length AR (AR-FL) mRNA has exons 1–8, whereas AR-V7 has exons 1–3 and cryptic exon 3 (exon 3b) (Hu et al., 2009) resulting in a version of AR that contains the *N*-terminal DNA-binding domains but lacks the ligand-binding domain (Fig 1A). AR-V7 can heterodimerize with AR-FL to activate canonical AR target genes in a ligand-independent manner (Watson et al., 2010). In addition, it can homodimerize with itself to constitutively activate a new set of gene targets that promote epithelial mesenchymal transition (EMT) and metastasis (Han et al., 2024). Because it lacks the ligand-binding domain, AR-V7 can circumvent the pharmacological effects of AR antagonists, contributing to the development of CRPC (Li et al., 2013). Developing novel therapies that overcome AR-V7 activity in CRPC remains clinically needed.

**Figure 1.**
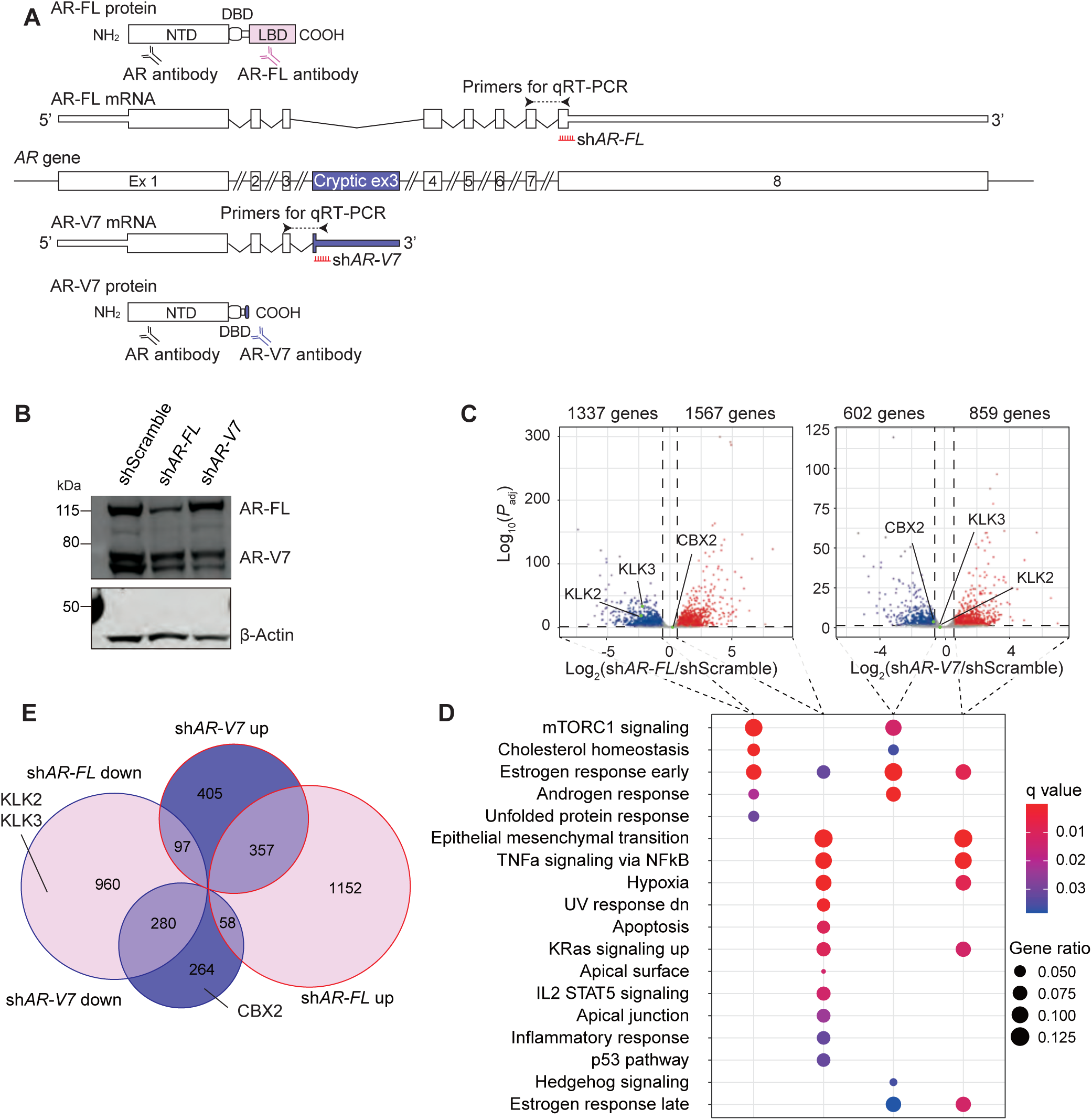
AR-FL- and/or AR-V7-targeted genes. (A) The gene structures of AR-FL and AR-V7. Antibodies, primers, and shRNAs used in this study are shown. AR protein levels (B) and RNA-seq (three independent experiments) (C–E) for 22Rv1 cells transduced with sh*AR-FL* or sh*AR-V7* and incubated in CSS-containing medium for 72 hr. (C) In the volcano plot, the genes whose expression was significantly decreased or increased (*P*_adj_ < 0.05 and FC > 1.5) by transduction with sh*AR-FL* or sh*AR-V7* are shown in blue or red, respectively. (D) The significantly enriched MSigDB_Hallmark gene sets in DEGs are shown. (E) Overlap of the DEGs with AR-FL or AR-V7 knockdown. Representative Western blots of three independent experiments are shown.

The switch/sucrose non-fermentable (SWI/SNF) is a large (1–2 MDa) evolutionarily conserved multi-subunit chromatin remodeling complex that regulates DNA accessibility through ATP-dependent nucleosome repositioning (Smith et al., 2003; Kadoch et al., 2016). In mammalian cells, three main types of SWI/SNF complexes have been described based on their distinct composition: canonical brahma-related gene 1 (BRG1)/brahma (BRM)-associated factor (cBAF), polybromo BAF (PBAF), and glioma tumor suppressor candidate region (GLTSCR) 1/1L-BAF (GBAF) (Mashtalir et al., 2018; Alpsoy et al., 2018). While these complexes share core components, such as the ATPase subunits BRG1 or BRM (SMARCA4/2) and structural subunits BAF155 and BAF170 (SMARCC1/2), they incorporate distinct accessory factors, including AT-rich interactive domain-containing protein (ARID) 1A/B and double PHD fingers (DPF)1/2/3 in cBAF, ARID2, polybromo-1 (PBRM1), and bromodomain containing (BRD) 7 in PBAF, and GLTSCR1/1L and BRD9 in GBAF (Wang et al., 1996; Kaeser et al., 2008; Alpsoy et al., 2018). The three SWI/SNF complexes show differential genomic binding patterns. cBAF is predominantly enriched at enhancers, PBAF is observed at promoters and gene bodies, and GBAF primarily localizes to promoters and CCCTC-binding factor (CTCF) sites (Michel et al., 2018). SWI/SNF complexes play central roles in transcriptional regulation across diverse biological processes.

SWI/SNF complexes have long been implicated in AR-mediated transcriptional regulation and prostate cancer progression (Ordonez-Rubiano et al., 2024). Previous studies, including our own, have shown that inhibition or degradation of several SWI/SNF components attenuates canonical AR signaling in both therapy-sensitive and CRPC (Alpsoy et al., 2021; Xiao et al., 2022; Shishodia et al., 2022; Ordonez-Rubiano et al., 2023). Given the central role of AR-V7 in CRPC and the difficulty of directly targeting it, identifying regulatory factors that control AR-V7 function may reveal actionable therapeutic vulnerabilities. Here we investigated the role of SWI/SNF complexes in regulating AR-V7 function. We found that cBAF is the only SWI/SNF subcomplex that regulates AR-V7-specific function and, unexpectedly, it does so by increasing the expression level of AR-V7, not its activity. Through mechanistic analysis we found that, in addition to alternative splicing, AR-V7 results from premature transcriptional termination. The premature transcriptional termination is facilitated by cBAF, which binds at an intronic region of AR downstream of cryptic exon 3. There it facilitates accessibility for enhancer activation and promoter looping that prevents the progression of polymerase. Inhibition or genetic deletion of cBAF subunits reduces cBAF binding, accessibility, and promoter looping at this site, resulting in transcriptional readthrough and restoration of the AR-FL transcript.

## Materials and Methods

### Chemicals and reagents

Dihydrotestosterone (DHT) was purchased from Merck (Frankfurter, Germany). ACBI-1, VZ-185, and BI-7273 were kindly provided by Boehringer Ingelheim (Ingelheim, Germany). BD98 was synthesized as described (Marian et al. 2018). iBRD9, JQ1, enzalutamide (ENZA), BRM014, and suberoylanilide hydroxamic acid (SAHA) were purchased from Cayman Chemical (Ann Arbor, MI). dBRD9 and AGB1 were obtained from Tocris (Minneapolis, MN). Doxycycline (Dox) was purchased from EMD Chemicals (San Diego, CA).

### Cell culture

Human prostate carcinoma-derived 22Rv1 (RRID: CVCL_1045) and LNCaP (RRID: CVCL_1379) cells (ATCC, Manassas, VA), were cultured in Roswell Park Memorial Institute (RPMI) 1640 supplemented with 10% fetal bovine serum (FBS) (Corning, Corning, NY), 10 mM 4-(2-hydroxyethyl)-1-piperazineethanesulfonic acid (HEPES) (Cytiva, Marlborough, MA), 1×minimum essential medium non-essential amino acids (MEM NEAA) (Corning), 100 U/mL penicillin and 100 μg/mL streptomycin (Merck), and 2 mM L-alanyl-L-glutamine (Corning), and 2.5 μg/mL plasmocin (Invivogen, San Diego, CA). Human prostate carcinoma-derived VCaP (RRID: CVCL_2235, ATCC) were cultured in Dulbecco’s Modified Eagle Medium/Nutrient Mixture F-12 (DMEM/F12) supplemented with 10% FBS, 10 mM HEPES, 1×MEM NEAA, 1 mM sodium pyruvate (Corning), 100 U/mL penicillin and 100 μg/mL streptomycin, and 2 mM L-alanyl-L-glutamine, and 2.5 μg/mL plasmocin. HEK293T cells (RRID: CVCL_0063, ATCC) were cultured in Dulbecco’s modified Eagle’s medium supplemented with 10% FBS, 1 mM sodium pyruvate, 100 U/mL penicillin and 100 μg/mL streptomycin, and 2 mM L-alanyl-L-glutamine, and 2.5 μg/mL plasmocin. For androgen deprivation studies, charcoal-stripped FBS (CSS) was used at 10% in phenol red-free RPMI 1640 or DMEM/F12.

### Lentiviral constructs

Short hairpin RNAs (shRNAs) targeting AR-FL, AR-V7, phosphatase 1 nuclear targeting subunit (PNUTS) (Supplemental Table 1) were cloned into pLKO.1-puro (Addgene plasmid #8453). GIPZ Lentiviral shRNAs (Horizon, Cambridge, UK) (Supplemental Table 2) were kindly provided by Dr. Diana Hargreaves (Salk Institute, CA).

The coding region of AR-V7 was amplified with an in-frame *C*-terminal FLAG tag by PCR and cloned into TetO-FUW-puro (Chowdhury et al., 2016, Addgene plasmid #85747) using In-Fusion HD Cloning Kit (Takara Bio, Kusatsu, Japan). pLenti CMV rtTA3 Hygro (w785-1) was a gift from Eric Campeau (Addgene plasmid #26730).

A lentiviral expression plasmid for TurboCas (dCas9-miniTurbo) was a gift from Ali Shilatifard (Cenik et al., 2024; Addgene plasmid # 232346). The blasticidin selection marker was replaced with the puromycin resistance gene (TurboCas-puro).

Lentiviral expression plasmids for dCas9-hemagglutinin (HA), MS2-FK506-binding protein (FKBP)-rapamycin binding domain of mTOR (FRB), and Dpf2-FKBP-V5 were previously established (Braun et al., 2017; Bergwell et al., 2024). sgRNAs targeting the *AR* gene (Supplemental Table 3) were cloned into lenti sgRNA(MS2) zeo backbone (Addgene plasmid #61427) or pLV-U6-gRNA-UbC-eGFP-P2A-Bsr, a gift from Charles Gersbach (Klann et al., 2017, Addgene plasmid #83925). lenti-EF1a-dCas9-VPR-Puro was a gift from Kristen Brennand (Ho et al., 2017, Addgene plasmid # 99373). Because Zukher et al. (2023) reported that dCas9 recruited by an sgRNA targeting the antisense strand can pause RNA polymerase II elongation, sgARint3#1 and sgARint3#2 were designed to target the sense strand.

### Lentiviral production

HEK293T cells were seeded in a 15-cm culture dish overnight and transfected with each lentiviral construct, along with the packaging vectors pMD2.G (Addgene plasmid #12259) and psPAX2 (Addgene plasmid #12260). After 16 hr of incubation, the medium was changed. The supernatant was collected 48 hr later by ultracentrifugation at 17,500g for 2 hr at 4 ℃. The pellet was then resuspended in 100 μL of phosphate-buffered saline (PBS) and stored at -80 ℃.

### Lentivirus infection

To establish stable knockdown systems, 22Rv1 cells were seeded in 12-well plates and incubated for 24 hours to reach 80% confluence, after which lentivirus encoding pLKO.1 or GIPZ shRNA was added. Twenty-four hours after transduction, the cells were selected by treatment with puromycin.

For inducible systems, 22Rv1 cells were first transduced with rtTA3 and selected with hygromycin, then transduced with TetO-AR-V7 constructs and selected with puromycin. Resulting cell lines were designated 22Rv1/Tet-Empty and 22Rv1/AR-V7-FLAG.

For establishment of TurboCas system, 22Rv1 cells were transduced with TurboCas-puro along with sgCtrl or sgARint3#1/2 and selected using puromycin and blasticidin. Resulting cell lines were designated 22Rv1/TurboCas/sgCtrl and 22Rv1/TurboCas/sgARint3.

To establish 22Rv1 cells with the Dpf2 recruitment system, the cells were sequentially transduced with dCas9-HA, MS2-FRB, and Dpf2-FKBP-V5 and selected using blasticidin, hygromycin, and puromycin (Supplemental Fig. 7B). The established cells were co-transduced with a sgRNA pair targeting each region and selected using Zeocin.

To establish the CRISPR activation (CRISPRa) system for the intron 3 of AR, 22Rv1 cells were transduced with dCas9-VP64-p65-Rta (VPR) along with sgCtrl or sgARint3#1/2 and selected using puromycin and blasticidin.

### ARID1B BromoTag (BT) system

To generate 22Rv1 cells expressing ARID1B-BT fused protein, C-terminal tagging of ARID1B with BT was conducted with CRISPR knock-in (Supplemental Fig. 4A). To construct a donor plasmid, we amplified the homology region around the stop codon of *ARID1B* (1127 bp) using ARID1B LHA S SacI and ARID1B RHA AS KpnI primers listed in Supplemental Table 4 and cloned it into pBluescript II. Subsequently, inverse PCR was performed using ARID1B LHA AS BamHI and ARID1B RHA S BamHI primers (Supplemental Table 4), and the BT tagging cassette, derived from pMK388 (Hatoyama et al., 2024, Addgene plasmid #121189), was inserted immediately upstream of the stop codon using the In-Fusion HD Cloning Kit. We obtained the linearized donor DNA by PCR using ARID1B LHA S and ARID1B RHA AS primers and transfected it into 22Rv1 cells using Lipofectamine 3000 (Thermo Fisher Scientific, Waltham, MA). After incubation for 24 hr, the cells were transfected with single-guide RNA (sgRNA) targeting around the stop codon of ARID1B (sgARID1B, Supplemental Table 3), synthesized by Integrated DNA Technologies (Coralville, IA), and Cas9 protein (Thermo Fisher Scientific) using Lipofectamine CRISPRMAX Cas9 (Thermo Fisher Scientific), and 1 μM Alt-R HDR Enhancer V2 (IDT) was added to promote the knock-in. After selecting the knocked-in cells with blasticidin, we performed single-cell cloning. By genotyping using primer pairs a (ARID1B ex 18 S and ARID1B 3’-UTR AS), b (ARID1B ex 18 S and BromoTag AS), and c (BSD S and ARID1B 3’-UTR AS), a clone with bi-allelic insertion was obtained. The resulting cells were designated 22Rv1/ARID1B-BT and transduced with sh*ARID1A*#1 as described above.

### Treatment of 22Rv1 and VCaP cells with compounds

22Rv1 cells were treated with 10 nM DHT for 16 hr in a CSS-containing medium. 22Rv1 (and its derivatives), LNCaP, or VCaP cells were treated with 0.5 μM ACBI-1, 5 μM BD98, 5 μM BI-7273, 0.5 μM VZ185, 5 μM iBRD9, 0.5 μM dBRD9, 1 μM JQ1, 10 μM ENZA, 5 μM BRM014, 200 ng/mL Dox, 5 nM rapamycin, 200 nM AGB1, or 5 μM SAHA in CSS- or FBS-containing medium.

### Partial deletion of the intron 3 of *AR*

To delete the intron 3 of *AR*, pX333 (Addgene plasmid #64073), which enables the expression of Cas9 and two sgRNAs, was used. Sequences of sgARint3#3 and sgARint3#4 targeting the intron 3 of *AR* (Supplemental Table 3) were cloned into pX333, and the construct was termed pX333/sgARint3×2.

22Rv1 cells were transfected with pX333/sgARint3×2 using Lipofectamine 3000 along with pLKO.1-puro in a molar ratio of 10: 1 to confer puromycin resistance. After the selection with puromycin for 72 hr, the genomic DNA was prepared according to a previously published paper (Koh, 2013). To evaluate the deletion efficiency, end-point PCR was performed using AR int3 S and AR int3 AS (Supplemental Table 5), and the PCR product was subjected to agarose gel electrophoresis.

### Transfection with antisense oligonucleotide (ASO)

Gapmer Antisense oligonucleotides for control and ARe (Supplemental Table 6) were synthesized in Integrated DNA Technologies. 22Rv1 cells were transfected with 100 nM AsO using Lipofectamine 3000.

### Preparation of whole cell lysates, cytoplasmic extract (CE), and nuclear extract (NE)

Whole cell lysates were prepared as follows. The cells were washed once with PBS buffer and then resuspended in RIPA buffer [50 mM Tris pH 8.0, 150 mM NaCl, 1% v/v NP40, 0.1% w/v sodium dodecyl sulfate (SDS), 0.5% w/v sodium deoxycholate]. A cocktail of protease inhibitors was added to the buffer right before use. The samples were then incubated for 20 min on ice and centrifuged at 15,000g for 30 min at 4 ℃, and the supernatant was collected.

CE and NE were prepared as follows. The cells were washed once with PBS buffer and then resuspended in buffer A [20 mM HEPES pH 7.9, 25 mM KCl, 10% glycerol, 0.1% NP-40 with protease inhibitors] at a concentration of 20 million cells/mL. The cells were kept on ice for 5 min, and nuclei were isolated by centrifugation at 600g for 10 min. After collecting the supernatant as cytoplasmic extract, the nuclear pellet was washed in buffer A without NP-40 lysis buffer. The pellet was resuspended in lysis buffer [20 mM HEPES pH 7.9, 150 mM NaCl, 7.5 mM MgCl_2_, 1% TritonX-100 with protease inhibitors] at a concentration of 200 million nuclei/mL and rotated at 4 °C for 30 min. The lysate was cleared by centrifugation at 21,000g for 30 min.

### Glycerol gradient sedimentation analysis

Glycerol gradient sedimentation was performed according to the published protocol (Alpsoy and Dykhuizen, 2018). 22Rv1 cells (∼40,000,000 cells) were harvested, lysed in buffer A [25 mM HEPES pH 7.6, 5 mM MgCl_2_, 25 mM KCl, 0.05 mM EDTA, 10% glycerol, 0.1% Nonidet P-40 (NP-40) with protease inhibitors], and washed once with buffer A without NP-40. The nuclei were resuspended in buffer C [10 mM HEPES pH 7.6, 3 mM MgCl_2_, 100 mM KCl, 0.1 mM EDTA, 10% glycerol with protease inhibitors]. Ammonium sulfate (0.3 M final) was added to the nuclear suspension and rotated at 4 °C for 30 min. The chromatin pellet was removed by ultracentrifugation at 150,000g for 30 min. Ammonium sulfate powder (0.3 g/mL) was added and incubated on ice for 20 min. Proteins were precipitated by ultracentrifugation at 150,000 × *g* for 30 min. The protein pellet was resuspended in 100 μL of HEMG1000 buffer [25 mM HEPES pH 7.6, 0.1 mM EDTA, 12.5 mM MgCl_2_, 100 mM KCl with protease inhibitors]. The protein extracts (0.8 mg) were layered over a 10 mL 10–30% glycerol gradient and fractionated by centrifugation at 40,000 rpm (XL-100K; Beckman Coulter, Brea, CA) for 16 hr using the SW32Ti rotor (Beckman Coulter). Twenty fractions of 0.5 mL were collected per sample and were subjected to Western blotting.

### Western blotting

The samples were prepared for Western blotting by mixing them with a 1:3 ratio with a mixture of β-mercaptoethanol and 4X Bolt LDS Sample Buffer (Thermo Fisher Scientific) in a 1:10 ratio, then boiled at 95 °C for 5 min. The denatured samples were separated by 4–12% SDS-polyacrylamide gels (Thermo Fisher Scientific) and transferred onto Immobilon-FL PVDF membranes (Merck). The membranes were then blocked for 1 hr in Immobilon Signal Enhancer (Merck), stained overnight with primary antibodies (Supplemental Table 7), and probed with the infrared dye-labeled goat anti-mouse or anti-rabbit antibodies (LI-COR, Lincoln, NE). Images were obtained using an Odyssey Clx imager (LI-COR).

### RT-qPCR and end-point RT-PCR

RNA was extracted using TRIzol (Thermo Fisher Scientific), and cDNA was synthesized using Verso cDNA Synthesis Kit (Thermo Fisher Scientific) with 3:1 oligo dT: random hexamer oligonucleotides. For the evaluation of the ARe level, ARe antisense primer was used as a primer. RT-qPCR was performed using PowerUp SYBR Green Master Mix (Thermo Fisher Scientific) in the CFX qPCR instrument (Bio-Rad, Hercules, CA). The primers are listed in Supplemental Table 7. End-point RT-PCR was conducted using primer pair of AR ex2 S and AR ex 5 AS or that of AR ex2 S and AR ex3b AS (Supplemental Table 5).

### RNA-seq

RNA was extracted using TRIzol and further purified by column purification with the PureLink RNA Mini Kit (Thermo Fisher Scientific) and on-column PureLink DNase (Thermo Fisher Scientific) treatment. DNA libraries were generated using the Illumina Stranded mRNA Prep, Ligation (Illumina, San Diego, CA) with sets of IDT for Illumina RNA UD Indexes (Illumina), and the concentration was determined using Bioanalyzer (Agilent Technologies, Santa Clara, CA). Libraries were pooled with equimolar amounts using cluster numbers obtained from MiSeq (Illumina) and sent for 150 bp paired-end sequencing on the NovaSeq 6000 or NovaSeq X platform (Novogen, Sacramento, CA).

Sequencing data were processed using Partek workflow. Read alignment to the human genome build hg38 was performed with STAR (Dobin et al., 2013). The resulting data for genes of interest were visualized using the Integrative Genomics Viewer (Robinson et al., 2011; Thorvaldsdottir et al., 2013). The number of reads aligned to genes was quantified with HTSeq. RNAseqChef (Etoh and Nakao, 2023) was used to assemble gene expression data, and differential gene expression analysis was conducted using DESeq2 (Love et al., 2014). The Benjamini-Hochberg method was used to adjust the *P*-values for multiple testing. Quantitative overlap analysis was performed using Eulerr (https://eulerr.co).

### Evaluation of cell viability

The cells were seeded in a white 96-well plate with clear bottom (Greiner Bio-One, Kremsmünster, Austria) and treated with compounds as described above. Cell viability was measured with a CellTiter-Glo kit (Promega, Madison, WI) in a GloMax Discover plate reader (Promega).

### Chromatin immunoprecipitation

22Rv1 cells were dual crosslinked with 2 mM disuccinimidyl glutarate (Covachem, Loves Park, IL) for 45 min at 25 °C, followed by 1% formaldehyde (Thermo Fisher Scientific) for 10 min at 25 °C, as described in Tian et al. (2012). Crosslinking was quenched with 125 mM glycine, for 10 min at 25 °C. The cells were washed with ice-cold PBS, scraped into 15-mL tubes, pelleted, flash-frozen in liquid nitrogen, and stored at -80 °C until ready for processing. Cell pellets were thawed on ice, extracted with nuclear extraction buffer [50 mM HEPES-KOH pH 8.0, 140 mM NaCl, 1 mM EDTA, 10% glycerol, 0.5% NP-40, 0.25% Triton X-100] for 10 min on ice, and washed once with nuclear wash buffer [10 mM Tris-HCl pH 8.0, 1 mM EDTA, 0.5 mM EGTA, 200 mM NaCl]. Nuclear pellets were resuspended in shearing buffer [10 mM Tris-HCl pH 8.0, 1 mM EDTA, 0.1% SDS] and sonicated using Branson SFX250 (Emerson, St. Louis, MO) at 10% amplitude for 8 min (ChIP-qPCR) or 16 min (ChIP-seq) at 0.5 sec ON and 1.5 sec OFF setting to obtain 200–1000 bp (ChIP-qPCR) or 100–500 bp (ChIP-seq) fragment size. The chromatin was quantified using absorbance at 260 nm and further diluted with 2×ChIP dilution buffer [100 mM HEPES-KOH pH 7.5, 600 mM NaCl, 1 mM EDTA, 2% Triton X-100, 0.2% sodium deoxycholate, 0.1% SDS] to obtain equal chromatin amounts (∼5,000,000 nuclei for ChIP-qPCR or ∼20,000,000 nuclei for ChIP-seq) in equal volume. After overnight incubation at 4 °C with rotation with the primary antibody (Supplemental Table 7), Protein A or Protein G Dynabeads (5 μL per μg of antibody) were added and rotated at 4 °C for another 2 hr. The beads were sequentially washed with low-salt wash buffer [20 mM HEPES-KOH pH 7.5, 0.1% SDS, 0.1% deoxycholate, 1% Triton X-100, 150 mM NaCl, 1 mM EDTA, 0.5 mM EGTA], high-salt wash buffer [20 mM HEPES-KOH pH 7.5, 0.1% SDS, 0.1% deoxycholate, 1% Triton X-100, 500 mM NaCl, 1 mM EDTA, 0.5 mM EGTA], LiCl wash buffer [20 mM HEPES-KOH pH 7.5, 0.5% deoxycholate, 0.5% IGEPAL CA-630, 250 mM LiCl, 1 mM EDTA, 0.5 mM EGTA], and final wash buffer [20 mM HEPES-KOH pH 7.5, 1 mM EDTA, 0.5 mM EGTA]. The immunoprecipitated chromatin was eluted once with 200 μL and once with 100 μL elution buffer [100 mM NaHCO_3_, 1% SDS] for 30 min each at 37 °C with shaking. The eluate and the saved input were treated with 20 μg RNase A (Thermo Fisher Scientific), 40 μg Proteinase K (Thermo Fisher Scientific) and reverse cross-linked at 65 °C for 16 hr. DNA was extracted once with phenol:chloroform; once with chloroform and precipitated by adding 1/10 volume of 3 M NaOAc pH 5.2; 1 volume 2-propanol and 40 μg glycogen (Thermo Fisher Scientific) for 1 hr at -80 °C. After centrifugation at top speed for 1 h, the pellet was washed with fresh 70% ethanol and then air-dried. DNA was resuspended in low-EDTA TE buffer.

qPCR was performed as mentioned above using the primers listed in Supplemental Table 9. ChIP-seq library was prepared with Ovation Ultralow System V2 UDI (Tecan Genomics, Redwood City, CA) according to manufacturer’s directions. The final library was subjected to double-sided selection using 0.65× and 1×SPRIselect (Beckman Coulter, Brea, CA) and sequenced by NovaSeq 6000 platform on 150bp PE sequencing.

### Cleavage under targets & release using nuclease (CUT&RUN)

CUT&RUN was performed using CUTANA ChIC/CUT&RUN Kit (Epicypher, Durham, NC). 22Rv1 cells (500,000 cells) were immobilized to concanavalin A beads, permeabilized with 0.05% digitonin, and treated with histone H3 lysine 27 acetylation (H3K27ac) antibody (Supplemental Table 7). After overnight incubation at 4 °C on a nutator, pAG-MNase was added, followed by calcium chloride to activate it and cleave the targeted DNA. After purification of the enriched DNA using SPRI beads, the sequencing library was prepared using CUTANA CUT&RUN Library Prep Kit (Epicypher) and sequenced by NovaSeq 6000 platform on 150bp PE sequencing.

### Assay for transposase-accessible chromatin (ATAC)-seq

ATAC-seq was processed following Corces et al. (2017). 22Rv1 cells (500,000 cells) were lysed in 50 μL Resuspension Buffer [10 mM Tris-HCl, pH 7.4, 10 mM NaCl, 3 mM MgCl_2_] supplemented with 0.1% NP-40, 0.1% Tween-20, and 0.01% digitonin and incubated on ice for 3 min. An additional 1 mL of RSB with 0.1% Tween-20 was added to each sample, and nuclei were pelleted by centrifugation at 600g for 10 min at 4 °C. The samples were then resuspended in 50 μL transposition mix [25 μL 2× TD buffer (Illumina), 2.5 μL transposase (100 nM final, Illumina), 16.5 μL PBS, 0.5 μL 1% digitonin, 0.5 μL 10% Tween-20, 5 μL H_2_O] and incubated in a thermomixer at 37 °C for 30 min with 1,000 rpm mixing. After transposition, DNA fragments were cleaned up using the MinElute PCR Purification Kit (Qiagen, Venlo, Netherlands) and PCR-amplified using indexed oligos and NEBNext High-Fidelity 2X PCR Master Mix (NEB, Ipswich, MA). The optimal number of amplification cycles for each sample was determined by qPCR. After cleanup with MinElute PCR Purification Kit, fragments between 100–1000bp were selected using AmpureXP beads (Beckman Coulter) and sequenced by AVITI (Element Biosciences, San Diego, CA) for 75 bp paired-end reads.

### Data analysis of ChIP-seq, ATAC-seq, and CUT&RUN

Raw paired-end sequencing reads were processed using Cutadapt (Martin, 2011) to remove adapter sequences and low-quality bases (Phred quality cutoff ≥ Q20, minimum read length 30 bp). The resulting reads were mapped to the human GRCh38 genome using Bowtie2 (Langmead and Salzberg, 2012) with the parameters --no-mixed and --no-discordant to ensure proper paired-end alignment. Using SAMtools (Li et al., 2009), mate information was corrected, duplicate reads were removed, reads with a mapping quality score ≥ 30 were retained, and mitochondrial reads were excluded. Reads overlapping with ENCODE blacklist regions (hg38-blacklist.v2) were filtered out with BEDTools (Quinlan and Hall, 2010). Genome-wide coverage tracks were generated using deepTools (Ramírez et al., 2016) with counts per million normalization. Peak calling was performed using MACS2 (Zhang et al., 2008) to identify enriched genomic regions with a p-value cutoff of 1 × 10^-4^. AR-FL-specific, AR-V7-specific, and AR-FL/AR-V7-shared peaks were identified using BEDTools intersect. We performed known motif discovery using the findMotifsGenome.pl program in HOMER (Heinz et al., 2010). To visualize the distribution of chromatin signal intensity around peak regions, we used deepTools. The signal intensity profiles from BigWig files were computed with the computeMatrix command, and the resulting matrix was visualized using the plotHeatmap and plotProfile commands.

### TurboCas

TurboCas was conducted according to a previously published paper (Cenik et al, 2024). 22Rv1 cells (∼80,000,000 cells) were treated with 500 *μ* M biotin for 30 min and lysed in lysis buffer [10 mM HEPES-KOH pH7.9, 10 mM KCl, 1.5 mM MgCl_2_, 340 mM sucrose, 10% glycerol, 0.2% Triton X-100 with protease inhibitors], followed by fractionation of nuclei in sucrose cushion buffer [10 mM HEPES-KOH pH7.9, 30% sucrose, 1.5 mM MgCl_2_]. The nuclear fraction was pelleted by centrifugation at 1,300g for 5 min at 4 °C, samples were digested at 4 °C for 45 min in chromatin buffer 1 [20 mM Tris-HCl pH 7.4, 150 mM NaCl, 1.5 mM MgCl_2_, 10% glycerol, 0.05% IGEPAL, 0.5 mM DTT, 250 U/ml Benzonase (Sigma, St. Louis, MO) with protease inhibitors]. After centrifugation at 20,000g for 10 min at 4 °C, the supernatant was collected as fraction 1, and the pellet was further solubilized by increasing salt concentration in chromatin buffer 2 [20 mM Tris-HCl pH 7.4, 500 mM NaCl, 1.5 mM MgCl_2_, 10% glycerol, 0.05% IGEPAL, 0.5 mM DTT, 3 mM EDTA with protease inhibitors] for 30 minutes. After centrifugation at 20,000g for 10 min at 4 °C, the supernatant (fraction 2) was combined with fraction 1. Western blotting for TurboCas and streptavidin staining were performed using HA antibody (Supplemental Table 7) and infrared dye-labeled streptavidin (LI-COR), respectively.

Following the protocol from (Hollenstein et al., 2023), streptavidin magnetic beads (Thermo Fisher Scientific) were acetylated with 5 mM sulfo-NHS-acetate by incubating for 60 min at room temperature to block cleavage of streptavidin by proteases in the following step. After washing three times in chromatin buffer 1, biotinylated proteins were purified using the biotinylated streptavidin beads by incubating overnight at 4 °C with gentle rotation. The next day, beads were washed twice in RIPA buffer [50 mM Tris pH 7.5, 150 mM NaCl, 0.1% SDS, 0.5% deoxycholate, 1% Triton X-100], once in 1 M KCl, once in 0.1 M Na_2_CO_3_, once in 2 M urea, two washes in RIPA buffer, and six washes in PBS at room temperature.

Peptides were analyzed on a timsTOF HT mass spectrometer (Bruker) coupled to an Evosep One LC system (Evosep). For each sample, 0.5 µg of peptides in 0.1% (v/v) formic acid was loaded onto Evotips Pure (Evosep) according to the manufacturer’s instructions and separated with the 60 samples-per-day method (24-min gradient) on a PepSep C18 column (8 cm × 150 µm i.d., 1.5 µm particles). The column was connected to a ZDV Sprayer with a 20 µm emitter in a CaptiveSpray source (Bruker). Mobile phase A was 0.1% (v/v) formic acid in water and mobile phase B was 0.1% (v/v) formic acid in acetonitrile.

Data were acquired in dia-PASEF mode. Each cycle consisted of one MS1 scan followed by 16 dia-PASEF scans, each containing two 25-m/z isolation windows (32 windows in total), covering m/z 400–1201 and 1/K0 0.6–1.6 V s cm-2 with 100 ms accumulation and ramp times. Collision energy increased linearly with ion mobility, from 20 eV at 1/K0 = 0.6 V s cm-2 to 59 eV at 1/K0 = 1.6 V s cm-2. Ion mobility was calibrated using three ions of the Agilent ESI-L Tuning Mix (m/z 622.0289, 922.0097 and 1221.9906). All samples were acquired with the same method.

Raw files were analyzed in Spectronaut (v19, Biognosys) using the library-free directDIA+ (Deep) workflow and the Pulsar search engine against the UniProt human reference proteome (UP000005640; downloaded 10/05/2025), supplemented with the TurboCas sequence and common contaminants. Trypsin/P was specified with up to 2 missed cleavages, peptide lengths of 7 – 52 residues and up to five variable modifications per peptide. Cysteine carbamidomethylation was set as a fixed modification, and methionine oxidation and protein N-terminal acetylation as variable modifications. Precursor, peptide and protein-group identifications were filtered at 1% FDR and cross-run normalization was enabled. Quantification was based on MS2 fragment-ion areas. Protein quantities were log2-transformed and normalized to TurboCas in each sample by subtracting its log2 intensity, to correct for differences in bait expression. Missing values were imputed in Perseus (Tyanova et al., 2016) from a normal distribution (width 0.3, downshift 1.8). The Benjamini-Hochberg method was used to adjust the P-values for multiple testing. Proteins which level was significantly enriched (*P*_adj_ < 0.05 and FC > 2) were subjected to an enrichment analysis using Enrichr (Xie et al., 2021) against the GO Biological Process 2026 library.

### Polymerase intact nascent transcript (POINT)-seq

POINT-seq was conducted according to a previously published paper (Sousa-Luís et al, 2021) with minor modifications. 22Rv1 cells (∼40,000,000 cells) were lysed with 4 mL of HLB+NC buffer [10 mM Tris-Cl pH 7.5, 10 mM NaCl, 2.5 mM MgCl_2_, 0.5% NP-40, 0.5% 3-((3-Cholamidopropyl) dimethylammonio)-1-propanesulfonate] on ice for 5 min. The lysate was underlaid with 1 mL of HLN+NS buffer [10 mM Tris-Cl pH 7.5, 10 mM NaCl, 2.5 mM MgCl_2_, 0.5% NP-40, 10% (w/v) sucrose]. The nuclear fraction was pelleted by centrifugation at 420g for 5 min at 4 °C and resuspended in 200 μL of ice-chilled NUN1 buffer [20 mM Tris-HCl pH 7.9, 75 mM NaCl, 0.5 mM EDTA, 50% glycerol] and then lysed in 2 mL of ice-cold NUN2E buffer [20 mM HEPES-KOH pH 7.6, 300 mM NaCl, 0.2 mM EDTA, 7.5 mM MgCl_2_, 1% NP-40, 1 M Urea, 5% Empigen BB, 1xProtease inhibitor Complete EDTA-free (Roche, Basel, Switzerland), 1xPhosSTOP (Roche)]. The chromatin was washed with 10 mL ice-cold PBS twice and then digested in DNase mixture [10 mM Tris-HCl pH 7.5, 400 mM NaCl, 10 mM MnCl_2_, 2 Units/μL RiboLock (Thermo Fisher Scientific), 0.2 Units/μL Turbo DNase (Thermo Fisher Scientific)] for 15 min at 37 °C with shaking. EDTA (final 1 mM) was added to stop the DNase reaction, and the supernatant was collected after centrifugation at 16,000g for 5 min at 4 °C. Ice-cold NET-2E buffer (800 µL) [50 mM Tris-HCl, pH 7.4, 150 mM NaCl, 0.05% NP-40, 5% Empigen BB] was used for the dilution of the supernatant. RPB1 antibody (10 μg) was conjugated with 200 μL anti-mouse IgG Dynabeads (Thermo Fisher Scientific) in 1 mL of NET-2 buffer [50 mM Tris-HCl pH 7.4, 150 mM NaCl, 0.05% NP-40] and incubated at 4 °C overnight. The conjugated beads were added to the supernatant and then incubated at 4°C for 1 h. After incubation, beads were washed 6 times with 1 mL of ice-cold NET-2E buffer. POINT fraction was purified using TRIzol and treated with Turbo DNase at 37 °C for 15 min. The POINTs were fragmented to 150–300 nucleotides for sequencing according to the protocol of NEBNext Ultra II Directional RNA library prep kit (NEB) and were sequenced by AVITI for 150 bp paired-end reads.

### Data analysis of POINT-seq

Trimmed reads were mapped to the human reference genome (hg38/GRCh38) using STAR with the following parameters: --outSAMmultNmax 1, --outMultimapperOrder Random, and --outSAMstrandField intronMotif. Sense- and antisense-strand-specific BAM files were generated using SAMtools. Strand-specific coverage files were generated using BEDTools genomecov and normalized to reads per million (RPM) and DESeq2 sizing factors based on gene-level counts obtained by featureCounts (Liao et al., 2014). Normalized bedGraph files were converted to BigWig format using bedGraphToBigWig. Replicate BigWig files were averaged using BEDTools unionbedg to generate mean coverage tracks for each condition.

Protein-coding genes were selected from GENCODE release 47 annotation based on the following criteria (Kopczyńska et al., 2026): (1) no overlap with other transcripts on the same strand, and (2) at least 6 kb distance from the nearest downstream gene on the same strand. Active genes were defined as protein-coding genes with a POINT-seq signal exceeding 1,000 counts in the non-treatment (NT) condition. To focus on genes whose transcriptional levels were not markedly changed, we further filtered for stable genes whose fold change of the POINT-seq signal in a gene body was less than 2-fold change. To assess transcriptional readthrough, a termination index (TI) was calculated for each gene as the ratio of the POINT-seq signal in the 6 kb downstream region of the annotated polyadenylation site (PAS) to the signal in the gene body. The change in TI upon BD98 treatment (ΔTI) was calculated as the ratio of TI in BD98-treated cells to TI in NT cells. Stable genes were classified into two groups: Class II (ΔTI > 2.0), representing genes with increased transcriptional readthrough by BD98 treatment, and Class I (ΔTI ≤ 2.0), representing genes without marked readthrough induction.

Metagene profiles were generated using deepTools. To generate strand-specific metagene profiles, plus-strand and minus-strand genes were processed separately using their respective sense and antisense BigWig files, and the resulting matrices were merged using computeMatrixOperations rbind. For antisense profiles, BigWig signals were inverted by multiplying by -1 before matrix computation. Metagene profiles were visualized using plotProfile.

### Formaldehyde-assisted isolation of regulatory elements (FAIRE) assay

FAIRE assay was conducted according to a published protocol (Simon et al., 2014). 22Rv1 cells (∼8,000,000 cells) were crosslinked with 1% formaldehyde for 5 min at 25 °C. Crosslinking was quenched with 125 mM glycine for 5 min at 25 °C. The cells were scraped into 15-mL tubes, washed with ice-cold PBS three times, flash-frozen in liquid nitrogen, and stored at -80 °C until ready for processing. Cell pellets were thawed on ice, resuspended in lysis buffer A [10 mM Tris-HCl pH 8.0, 100 mM NaCl, 1 mM EDTA, 1% SDS, 2% Triton X-100], and sonicated using Branson SFX250 at 20% amplitude for 20 min at 0.5 sec ON and 1.5 sec OFF setting to obtain 200–1000 bp fragment size. By treatment with 10 μg RNaseA for 30 min each at 37 °C, followed by decrosslinking with 20 μg proteinase K at 65 °C for 16 hr, input DNA was prepared. FAIRE DNA and input DNA were extracted once with phenol:chloroform, once with chloroform, and precipitated by adding 1/10 volume of 3 M NaOAc pH 5.2, 1 volume 2-propanol, and 40 μg glycogen (Thermo Fisher Scientific). After centrifugation at 12,000g for 15 min, the pellet was washed with fresh 70% ethanol and then air-dried. DNA was resuspended in Tris-HCl (pH 7.4). FAIRE DNA was treated with 10 μg RNaseA for 30 min each at 37 °C followed by 20 μg proteinase K at 65 °C for 16 hr and extracted using NucleoSpin Gel and PCR Clean-up (Takara). qPCR was performed as mentioned above using the primers listed in Supplemental Table 9.

### Chromosome conformation capture (3C) assay

3C assay was conducted according to a published protocol (Hagège et al., 2007). 22Rv1 cells (2,000,000 cells) were crosslinked with 2% formaldehyde for 10 min at 25 °C. Crosslinking was quenched with 125 mM glycine for 5 min at 25 °C. Cell pellets were lysed in 1 mL of ice-cold lysis buffer [10 mM Tris-HCl pH 7.5, 10 mM NaCl, 5 mM MgCl_2_, 0.1 mM EGTA] and incubated for 10 min. The pelleted nuclei after centrifugation at 400g for 5 min at 4 °C were resuspended in 100 μL of 1.2x FastDigest Green Buffer (Thermo Fisher Scientific), and 1.5 μL of 20% SDS was added. After incubation for 1 hr at 37 °C with shaking, 10 μL of 20% (v/v) Triton X-100 was added. After additional incubation for 1 hr at 37 °C with shaking, 8 μL of FastDigest BamHI (Thermo Fisher Scientific) was added, and the samples were incubated overnight at 37 °C with shaking. Eight μL of 20% SDS was added to the digested nuclei and incubated for 20 min at 65 °C with shaking. After adding 1.225 mL of 1.15x ligation buffer (Promega), 75 μL of 20% Triton X-100, the samples were incubated for 1 hr at 37 °C with shaking. After adding 20 U ligase (Promega), the samples were incubated for 4 hr at 16 °C followed by 30 min at room temperature. After adding 12μg of proteinase K, the ligated samples were de-crosslinked by incubation at 65 °C. After treating with 12 μg RNase A, the ligated DNA was extracted with phenol:chloroform and resuspended in 10 mM Tris-HCl pH 7.5. qPCR was performed as mentioned above using the primers listed in Supplemental Table 10.

### Evaluation of the cleavage of the 3’-end of AR-V7 pre-mRNA

To evaluate the cleavage efficiency of the 3’-end of AR-V7 pre-mRNA, chromatin-associated RNA was obtained through chromatin isolation as described in POINT-seq sample preparation, followed by RNA extraction using TRIzol. After cDNA was synthesized using AR-V7 DS AS primer, RT-qPCR was performed as described above using the primers listed in Supplemental Table 11.

## Statistical Analysis

A comparison of two groups was made with an unpaired, two-tailed Welch’s *t* test. Data from multiple groups were compared using analysis of variance followed by Tukey’s test or Dunnett’s test. A value of *P* < 0.05 was considered statistically significant.

## Results

### SWI/SNF complexes regulate AR-FL and AR-V7-mediated transcription

To establish their respective gene targets, we knocked down AR-FL or AR-V7 in the human CRPC cell line 22Rv1 and performed RNA-Seq. The analysis indicated that kallikrein related peptidases (*KLK2* and *KLK3)* were decreased upon knockdown of AR-FL but not AR-V7, and chromobox 2 (*CBX2)* was decreased upon knockdown of AR-V7 but not AR-FL (Fig. 1B–E). To identify the individual SWI/SNF subcomplexes that regulate AR-FL- and/or AR-V7-mediated transcriptional regulation we treated 22Rv1 cells with a variety of pharmacological inhibitors of SWI/SNF including ACBI-1 (SMARCA2/4 degrader, Farnaby et al., 2019), BRM014 (SMARCA2/4 inhibitor, Papillon et al., 2018), BD98 (ARID1A/cBAF inhibitor, Marian et al., 2018), BI-7273 (BRD7/9 co-inhibitor, Martin et al., 2016), VZ185 (BRD7/9 co-degrader, Zoppi et al., 2019), iBRD9 (BRD9 inhibitor, Theodoulou et al., 2016), and dBRD9 (BRD9 degrader, Remillard et al., 2017). We also treated cells with BRD4 inhibitor JQ1 (Filippakopoulos et al., 2010) which should inhibit all AR targets, or ENZA (AR antagonist, Tran et al., 2009), which should not inhibit AR targets in these cells (Fig. 2A). *KLK2* and *KLK3* mRNA levels are reduced by all SWI/SNF inhibitors, supporting previous studies showing that all three SWI/SNF complexes enhance canonical AR signaling (Fig. 2B) (Alpsoy et al., 2021; Xiao et al., 2022; Ordonez-Rubiano et al., 2023). In contrast, *CBX2* mRNA level was decreased only by ACBI-1, BRM014, or BD98, suggesting that cBAF selectively regulates the expression of AR-V7-targeted genes (Fig. 2B). In a cell viability assay, ACBI-1, BD98, iBRD9, and JQ1 exerted the greatest anti-proliferation effects on 22Rv1 cells (Fig. 2C). These results obtained with normal FBS medium were consistent with results obtained with charcoal stripped serum (CSS) without androgens (Supplemental Fig. 1A and B).

**Figure 2.**
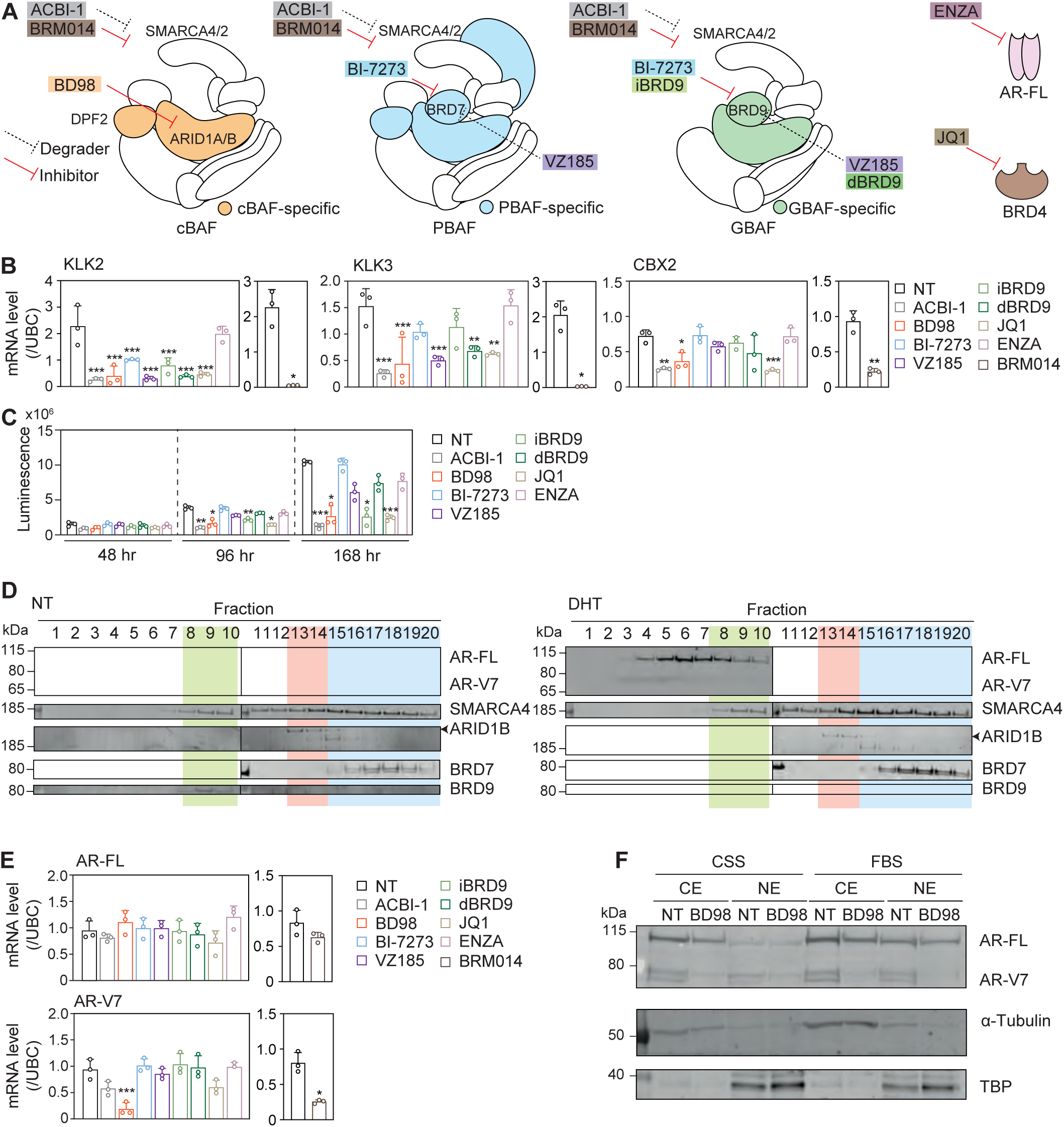
Effects of SWI/SNF inhibitors on AR-FL- and AR-V7-mediated transcriptional regulation and their expression. (A) Inhibitors targeting SWI/SNF components, AR-FL, and BRD4. KLK2, KLK3, CBX2 (B), AR-FL, and AR-V7 (E) mRNA levels, viability (C), and AR protein level in the CE and the NE (F) of 22Rv1 cells treated with 0.5 μM ACBI-1, 5 μM BD98, 5 μM BI-7273, 0.5 μM VZ-185, 5 μM iBRD9, 0.5 μM dBRD9, 5 μM JQ1, 10 μM ENZA, or 5 *μ* M BRM014 in FBS-containing medium for 48 (C), 96 (B, C, and E), or 168 hr (C). (D) Glycerol gradient sedimentation analysis for 22Rv1 cells treated with 10 nM DHT in CSS-containing medium for 16 hr. Representative Western blots of three independent experiments are shown. mRNA levels are relative to those in CSS-containing medium (Supplemental Fig 1A and D). Each column represents the mean ± SD of three independent experiments. \**P* < 0.05, \*\**P* < 0.01, and \*\*\**P* < 0.001, compared with NT.

Previous studies have established that SWI/SNF directly interacts with AR-FL (Li et al., 2006). To investigate whether SWI/SNF complexes interact with AR-V7 as well, a glycerol gradient sedimentation analysis was performed using 22Rv1 cells. Consistent with a previous report (Alpsoy and Dykhuizen, 2018), GBAF-specific BRD9, cBAF-specific ARID1B, and PBAF-specific BRD7, were detected in fractions 8–10, 13–14, and 15–20, respectively, with common subunit SMARCA4, which was detected in fractions 8–20 (Fig. 2D). AR-FL was broadly observed in fractions 3–20, and the band intensity was enhanced by DHT (Fig. 2D), a ligand of AR, suggesting that activated AR-FL interacts with the SWI/SNF complex. This is in agreement with reports that AR interacts with core subunits shared amongst SWI/SNF complexes, such as SMARCD1 (Van de Wijngaart, 2009), SMARCE1 (Link et al., 2005), and SMARCC1 (Hong et al., 2005). In contrast, AR-V7 was detected in fractions 2–7 but not with SWI/SNF complexes in fractions 8–20, in either the absence or presence of DHT (Fig. 2D). This supports the hypothesis that SWI/SNF interacts with AR via the hormone-induced coactivator binding groove in the ligand-binding domain (Van de Wijngaart, 2009), which is absent in AR-V7.

Since cBAF doesn’t act as a direct co-activator of AR-V7, we next checked whether SWI/SNF inhibitors regulate AR-FL and AR-V7 expression. Unexpectedly, treatment with ACBI-1 or BD98 decreased mRNA and protein levels of AR-V7 but not AR-FL (Fig. 2E, F, and Supplemental Fig. 1C). In VCaP cells, another CRPC cell line, treatment with BD98 also decreased AR-V7 levels and viability (Supplemental Fig. 1D–F). These results indicate that cBAF is required for the transcription of *AR-V7*.

### AR-V7 rescue of BD98-induced changes in gene expression and viability

To investigate how the cBAF regulation of AR-V7 expression affects the transcriptome separately from cBAF regulation of AR-FL activity, we established 22Rv1 cells with a tet-on-regulated AR-V7 expression system. Although there was some expression at baseline, treatment with doxycycline induced additional overexpression of AR-V7 in 22Rv1 cells (Fig. 3A). We performed RNA-seq analysis using 22Rv1/Tet-Empty and 22Rv1/Tet-AR-V7-FLAG cells, with and without BD98 treatment. In 22Rv1/Tet-Empty cells with BD98, mRNA levels of 2,564 genes, including KLK2, KLK3, and CBX2, were decreased, while mRNA levels of 2,056 genes were increased (Supplemental Fig. 2A). These differentially expressed genes (DEGs) partially overlapped with the genes whose levels were changed by knockdown of AR-FL or AR-V7 (Supplemental Fig. 2B), supporting that cBAF can modulate the expression of AR-FL and AR-V7-regulated genes. As shown in Fig. 3B, overexpression of AR-V7 partially mitigates the BD98-induced changes in gene expression (Clusters 1 and 3), including CBX2 but not KLK2 and KLK3. We identified eight significantly enriched MSigDB_Hallmark gene sets in the AR-V7-rescued expressed genes, primarily related to cell proliferation or inflammation (Fig. 3C). In agreement with the pathway analysis, exogenous AR-V7 partially restored the proliferation of 22Rv1 cells after BD98 treatment (Fig. 3D). These results indicate that BD98 exerts an anti-proliferative effect on 22Rv1 CRPC cells, in part by reducing AR-V7 expression.

**Figure 3.**
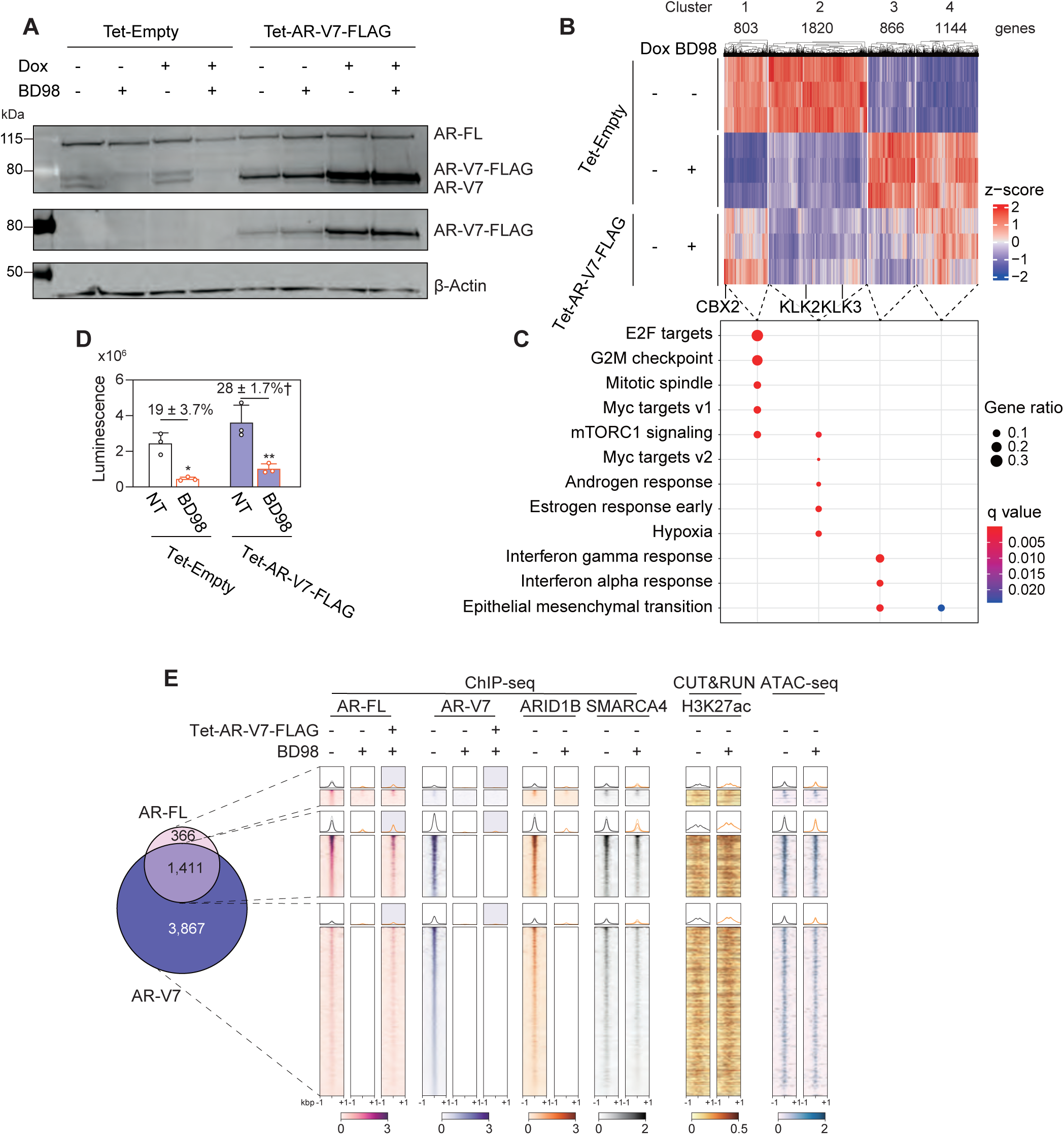
Effects of AR-V7 overexpression on BD98-mediated changes in gene expression. (A) AR protein levels of 22Rv1/Tet-Empty and 22Rv1/Tet-AR-V7-FLAG cells treated with 5 μM BD98 and/or 200 ng/mL Dox in CSS-containing medium for 96 hr. Representative Western blots of three independent experiments are shown. (B) RNA-seq for 22Rv1/Tet-Empty and 22Rv1/Tet-AR-V7-FLAG cells treated with 5 μM BD98 in CSS-containing medium for 96 hr (three independent experiments). k-means clustering was performed for DEGs (*P*_adj_ < 0.05 and FC > 1.5) with BD98 in 22Rv1/Tet-Empty cells. (C) The significantly enriched MSigDB_Hallmark gene sets are represented in each cluster. (D) The viability of 22Rv1/Tet-Empty and Tet-AR-V7-FLAG cells treated with 5 μM BD98 in CSS-containing medium for 168 hr. Each column represents the mean ± SD of three independent experiments. \**P* < 0.05 and \*\**P* < 0.01, compared with NT; †*P* < 0.05, compared with Tet-Empty. (E) ChIP-seq of AR-FL and AR-V7 for 22Rv1/Tet-Empty and 22Rv1/Tet-AR-V7-FLAG cells treated with 5 μM BD98 in CSS-containing medium for 48 hr (two independent experiments) and ChIP-seq of ARID1B and SMARCA4 (two independent experiments), CUT&RUN for H3K27ac (two independent experiments), and ATAC-seq (three independent experiments) for 22Rv1 cells treated with 5 μM BD98 for 48 hr in FBS-containing medium. Metagene plots (faint line, each replicate; dark line, mean of replicates) and heatmaps (mean of replicates) are shown for AR-FL-specific, AR-V7-specific, and their common regions.

Consistent with the established role for cBAF as a co-factor for AR-FL, the androgen response-related genes were decreased by BD98 and not rescued by the overexpression of AR-V7 (Fig. 3B). To validate that BD98 treatment reduces the binding of AR-FL to its target regions, we conducted ChIP-seq, CUT&RUN, and ATAC-seq. After treatment with BD98, ARID1B and SMARCA4 peak signals, which overlapped with H3K27ac signals, were reduced, resulting in a slight decrease in chromatin accessibility (Supplemental Fig. 2C). From the ChIP-seq using AR-FL and AR-V7-specific antibodies, 1,777 and 5,278 peaks were detected and were categorized into AR-FL-specific (366), AR-V7-specific (3,867), and shared (1,411) peaks (Fig. 3E). Motif analysis confirmed that these regions are enriched for consensus sequences for AR and forkhead box A1 (FOXA1), an AR-interacting pioneer factor (Supplemental Fig. 2D). In AR-FL-specific regions, AR-FL was colocalized with ARID1B/SMARCA4, and its binding to chromatin was decreased by treatment with BD98, in agreement with cBAF as a co-regulator of AR-FL. In AR-V7-specific and shared regions, AR-V7 peaks were eliminated with BD98 treatment. The signal was not restored by AR-V7 overexpression, most likely due to interference from the C terminal FLAG-tag; however, AR-FL peaks were restored with AR-V7 overexpression, suggesting that decreased AR-V7 expression by BD98 contributes to the reduction of AR-FL binding to the genome. While ARID1B/SMARCA4 signals were highest in the shared AR-FL/AR-V7 peaks, there was also enrichment in the AR-V7-specific regions, indicating that although cBAF doesn’t directly interact with AR-V7, it may still contribute to accessibility for AR-V7 DNA binding.

### The effects of ARID1A or ARID1B knockdown on AR-V7 expression

ARID1A and ARID1B are mutually exclusive components of cBAF (Wu and Roberts, 2013). While BD98 was developed as an ARID1A inhibitor (Marian et al., 2018), it most likely inhibits ARID1B as well as ARID1A, as is evident from the ChIP-Seq (Fig 3E). Thus, we examined whether ARID1A or ARID1B is more important for AR-V7 expression using RNA-Seq. Using lentiviral-mediated shRNA, ARID1A and ARID1B were successfully knocked down in 22Rv1 cells (Supplemental Fig. 3A and B). ARID1B and ARID1A protein levels were dramatically and slightly increased by knockdown of ARID1A and ARID1B, respectively, without changes in their mRNA levels, suggesting that ARID1A and ARID1B protein post-translationally compete for incorporation in the cBAF complex and can compensate for one another (Supplemental Fig. 3A and B). With regards to AR gene expression, sh*ARID1B* but not sh*ARID1A* decreased AR-V7 mRNA and protein levels (Supplemental Fig. 3B and C) and had a greater effect on 22Rv1 proliferation (Supplemental Fig. 3D); however, these effects were milder than global cBAF inhibition. Knockdown of ARID1B resulted in larger expression changes than knockdown of ARID1A (Supplemental Fig. 3E and F), most likely reflecting its greater abundance in 22Rv1 cells. In agreement with unique functions for ARID1A and ARID1B, many of the DEGs were unique for one or the other (Supplemental Fig. 3G). Many were even antagonistic; EMT pathway was positively enriched with sh*ARID1A* and negatively enriched with sh*ARID1B* perhaps reflecting the paralog compensation that occurs in the knockdowns (Supplemental Fig. 3B). In addition, many genes changed with cBAF inhibition are not affected in either sh*ARID1A* or sh*ARID1B*, indicating functionally redundancy at some genes.

To address this possibility, we employed the BT degron system (Bond et al., 2021) for acute depletion of ARID1B protein (Fig. 4A). Using CRISPR knock-in, we obtained 22Rv1 cells in which the ARID1B C-terminus is tagged with BT (22Rv1/BT cells, Supplemental Fig. 4A and B). The ARID1B band was shifted to a higher molecular weight by the knock-in and depleted by treatment with AGB1, a BT ligand, within 4 hr (Fig. 4B). Upon acute degradation of ARID1B there was no increase in ARID1A (Fig. 4B) and AR-V7 mRNA and protein levels were clearly decreased by 24 and 48 hr AGB1 treatment (Fig. 4B and C). RNA-seq analysis of AGB1-treated samples for 4 hr indicated that cBAF-regulated genes are classified into three categories: those concordantly regulated by both ARID1A and ARID1B (Clusters 1 and 6), those discordantly regulated by the two subunits (Clusters 3 and 7), and those regulated by only one of them (Clusters 2, 4, 5, and 8) (Fig. 4D and E). The effects of knockdown of ARID1B were expanded from 4 hr to 48 hr, and the DEGs were overlapped with those by BD98 treatment (Supplemental Fig. 4C). The pathways enriched among genes downregulated by ARID1A/ARID1B double knockdown, including CBX2 (Clusters 1 and 2, Fig. 4F and G), overlapped with those enriched among genes downregulated by BD98 and among genes enriched by AR-V7 overexpression (Clusters 1 and 2, Fig. 3B and C). These included cell proliferation-related and glycolysis- and hypoxia-related pathways. While the DEGs partially overlapped with those by AR-FL or AR-V7 knockdown in the same direction (Supplemental Fig. 4D), the androgen response pathway was enriched in the upregulated genes by ARID1B knockdown, possibly due to an increase in EP300 level (Fig. 4E and G), which has been reported to compensate for ARID1A loss (Wilson et al., 2020; Blümli et al., 2021) and to enhance AR signaling (Fu et al., 2000; Gioukaki et al., 2023) Collectively, cBAF would support aggressiveness of 22Rv1cells by regulating expression of a wide range of genes, including AR-V7.

**Figure 4.**
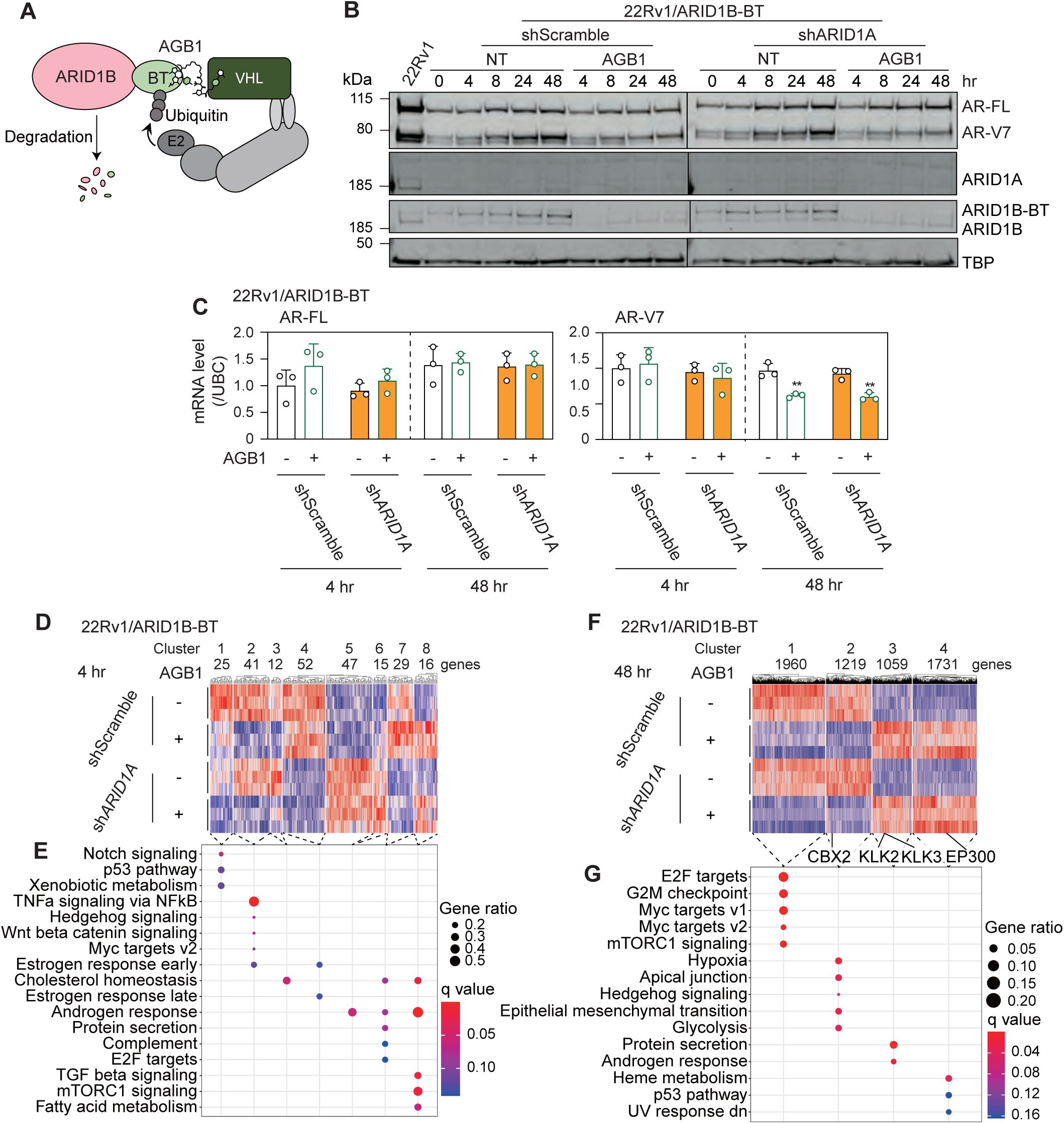
Effects of acute knockdown of ARID1B. (A) ARID1B-BT system. AR, ARID1A, and ARID1B protein (A) and AR-FL and AR-V7 mRNA (B) levels and RNA-seq (three independent experiments) (D–G) for 22Rv1/ARID1B-BT cells transduced with sh*ARID1A* and treated with 200 nM AGB1 for 4 (B–E), 8 (B), 24 (B), or 48 (B, C, F, G) hr in CSS-containing medium. Representative Western blots of three independent experiments are shown. Each column represents the mean ± SD of three independent experiments. \*\**P* < 0.01, compared with NT. k-means clustering (D and F) and the significantly enriched MSigDB_Hallmark gene sets (E and G) are represented.

### cBAF-mediated chromatin remodeling of the region downstream of cryptic exon 3

AR-V7 arises because of an alternative splicing event that includes a cryptic exon 3 that ends in transcriptional termination. Consistent with this, the RNA-seq coverage of cryptic exon 3 decreased following BD98 treatment in 22Rv1 cells, as well as following 48 h knockdown of ARID1B in 22Rv1/ARID1B-BT cells (Supplemental Fig. 6A). Jumonji domain containing 1A (JMJD1A), a histone demethylase, binds to the 3’ splice site in front of the cryptic exon 3 and recruits heterogeneous nuclear ribonucleoprotein F (HNRNPF), a splicing factor, to promote maturation of AR-V7 precursor mRNA (pre-mRNA) (Fan et al., 2018). Because JMJD1A interacts with ARID1A (Abe et al., 2015) and ARID1A/B can regulate splicing by facilitating nuclear speckles (Reddy et al., 2023), we hypothesized that cBAF also binds to the junction between intron 3 and cryptic exon 3 to facilitate JMJD1A binding to chromatin. However, ChIP-qPCR failed to identify SMARCA4 and ARID1A/B binding at the splice junctions in 22Rv1 cells (Supplemental Fig. 5A and B). Instead, we analyzed publicly available ChIP-seq data and noticed that SMARCA4 and ARID1A, but not BRD7 and BRD9, bind downstream of cryptic exon 3 in intron 3 (Supplemental Fig. 6A). ChIP-seq (Fig. 5A) and ChIP-qPCR (Supplemental Fig. 5C) in 22Rv1 cells confirmed that SMARCA4 and ARID1B bind to this region and that the enrichment was decreased by treatment with BD98. According to public ChIP-seq data, there is a peak of ATAC signal in AR-V7-positive 22Rv1 cells but not in AR-V7-negative LNCaP cells (Supplemental Fig. 6A). To test whether accessibility at this region is dependent on cBAF, we performed ATAC-seq (Fig. 5A) and FAIRE-qPCR (Supplemental Fig. 5D) with BD98 treatment and found that accessibility was indeed reduced with cBAF inhibition. Additionally, BD98 treatment decreased the level of H3K27Aac at this region (Supplemental Fig. 6D), consistent with reports of a dependency on cBAF for maintenance of H3K27ac at sites of active gene regulatory regions (Kelso et al., 2017). These results suggest that cBAF localizes downstream of cryptic exon 3 to maintain chromatin accessibility and histone acetylation in 22Rv1 cells.

**Figure 5.**
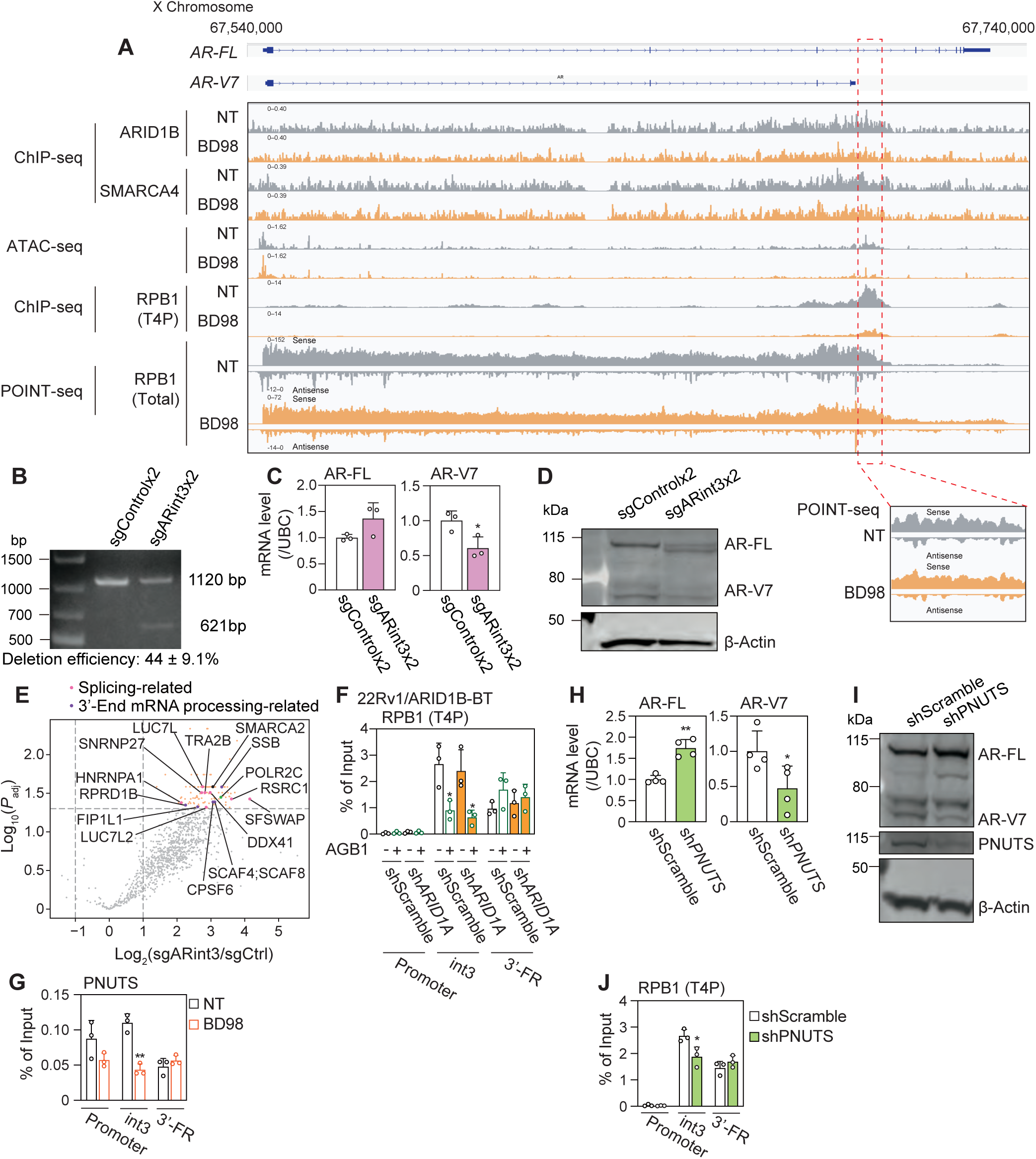
Effects of BD98 treatment on transcriptional termination of AR-V7. (A) Genomic tracks of ChIP-seq (ARID1B, SMARCA4, RPB1 (T4P)), ATAC-seq, and POINT-seq at the *AR* gene in 22Rv1 cells treated with 5 μM BD98 in FBS-containing medium for 48 hr. Each area represents the mean of two (ChIP-seq and POINT-seq) or three (ATAC-seq) independent experiments. (B–C) 22Rv1 cells were transfected with pX333/sgARint3×2 in FBS-containing medium for 72 hr. (B) The deletion was checked by end-point PCR. AR-FL and AR-V7 mRNA (C) and protein (D) levels. Representative Western blots and agarose gel electrophoresis of three independent experiments are shown. Each column represents the mean ± SD of three or four independent experiments. \**P* < 0.05, compared with sgControlx2. (E) TurboCas for 22Rv1/TurboCas/sgRNA. In the volcano plot, the proteins that were significantly enriched (*P*_adj_ < 0.05 and FC > 2) by transduction with sgARint3 are shown in orange. (F) ChIP-qPCR of RPB1 (T4P) in 22Rv1/ARID1B-BT cells transduced with shARID1A and treated with 200 nM AGB1 for 48 hr in CSS-containing medium. (G) ChIP-qPCR of PNUTS for 22Rv1 cells treated with 5 μM BD98 in FBS-containing medium for 48 hr. \**P* < 0.05 and \*\**P* < 0.01, compared with NT. AR and PNUTS mRNA (H) and protein (I) levels and ChIP-qPCR of RPB1 (T4P) (J) in 22Rv1 cells transduced with sh*PNUTS*. \**P* < 0.05 and \*\**P* < 0.01, compared with shScramble.

### The downstream region of cryptic exon 3 is important for AR-V7 expression

To examine whether the region downstream of cryptic exon 3 is required for AR-V7 expression, we deleted this region using the CRISPR-Cas9 system and two sgRNA targeting the intron 3 of *AR* (Fig. 5B). Confirming the importance of this region, AR-V7 mRNA and protein levels were decreased (Fig. 5C). Surprisingly, in Western blotting, a new additional band was observed below AR-FL (Fig. 5D), indicating that the CRISPR-mediated deletion might have caused exon skipping. We performed end-point RT-PCR using primers designed to cross exons 2 and 5 and observed a band at 700 bp in the control cells and new bands at 550 and 450 in the sgARint3 cells (Supplemental Fig. 6B). Since the genomic region around exon 3 is tandemly duplicated in 22Rv1 cells (Supplemental Fig. 6A) the two bands represent the loss of one or both copies of exon 3 (Supplemental Fig. 6B). While AR-V7 is significantly reduced in the sgARint3 cells (Fig. 5C), splicing of AR-V7 was not changed (Supplemental Fig. 6C). Collectively, this indicates that the region downstream of cryptic exon 3 is crucial for production of AR-V7 mRNA as well as the inclusion of exon 3 in AR-FL.

### Regulation of AR-V7 expression by transcriptional termination

To comprehensively identify proteins binding to the downstream region of AR-V7, we used TurboCas, a method that combines a proximity-labeling enzyme with CRISPR-dCas9 to enable region-specific labeling of chromatin factors (Cenik et al., 2024). By recruiting TurboCas to the downstream region of cryptic exon 3 followed by biotin treatment and streptavidin purification (Supplemental Fig. 7A), 112 proteins, including SMARCA2, were identified (Supplemental Table 12). Enrichment analysis revealed significant overrepresentation of splicing factor-related pathways (Supplemental Table 13). These factors may be involved in the inclusion of exon 3 in AR-FL. In addition, 3’-End mRNA processing factors were included, as well as POLR2C, an RNA polymerase II subunit. This result prompted us to focus on transcriptional termination.

During transcriptional termination RNA polymerase II passes the polyadenylation signal at the 3’ end of the gene and cleavage and polyadenylation factors are recruited to cleave the newly synthesized pre-mRNA and pause RNA polymerase II (Proudfoot, 2016). In 22Rv1, there is a RNA polymerase II peak downstream of cryptic exon 3 that is absent in AR-V7-negative LNCaP cells (Supplemental Fig. 6A), prompting us to examine whether cBAF binding at this region is important for transcriptional termination of AR-V7 mRNA. Upon treatment of cells with BD98 there was a reduction in the enrichment of RNA polymerase II subunit B1 (RPB1) and cleavage and polyadenylation specificity factor (CPSF) 73 in the transcriptional termination region of AR-V7, but not in the transcriptional termination region of AR-FL (Supplemental Fig. 5E). This indicates that cBAF is required for premature termination for *AR-V7* but isn’t required for all transcriptional termination. To confirm this, we performed ChIP-Seq of RPB1 phosphorylated at threonine 4 (T4) in the *C*-terminal domain (CTD), a mark associated with transcriptional termination (Kopczyńska et al., 2025). Indeed, RPB1 T4P enrichment downstream of cryptic exon 3 was decreased by treatment with BD98 in 22Rv1 cells (Fig. 5A) and by ARID1B knockdown in 22Rv1/ARID1B-BT cells (Fig. 5F).

Another regulator of transcriptional termination is the protein phosphatase (PP) 1-PNUTS complex, which decelerates RNA polymerase II in transcriptional termination regions (Cortazar et al., 2019) and has been reported to colocalize with SMARCA4 (Verheyen et al., 2015). Treatment with BD98 also reduces enrichment of PNUTS downstream of cryptic exon 3 (Fig. 5G), but not at the transcription termination site of *AR-FL*. Knockdown of PNUTS also reduces T4P level in the transcriptional termination region and a reduction in AR-V7 mRNA and protein (Fig. 5H–J).

Confirming a role for cBAF in transcriptional termination, we observe an increase in transcriptional readthrough downstream of cryptic exon 3 with BD98 treatment using POINT-seq, a nascent transcriptomics method (Fig. 5A). Collectively, these data indicate that cBAF is important for the transcriptional termination of *AR-V7*.

### Regulation of transcriptional termination of *AR-V7* by the chromatin environment

To determine whether the decreased H3K27ac level observed with BD98 treatment (Supplemental Fig. 5D) contributes to reduced AR-V7 levels, 22Rv1 cells were treated with SAHA, a histone deacetylase inhibitor. Treatment with BD98 decreased total H3K27ac level, which was restored upon co-treatment with SAHA (Supplemental Fig. 8B). SAHA also partially restored reduced RPB1 enrichment and reduced AR-V7 levels induced by BD98 (Supplemental Fig. 8A–C). In LNCaP cells, SAHA treatment increased *AR-V7* mRNA level but not *AR-FL* (Supplemental Fig. 8D), further confirming the importance of acetylation in premature transcriptional termination for AR-V7 expression.

To evaluate the contribution of cBAF-mediated chromatin accessibility to AR-V7 expression, we established an inducible recruitment system using Dpf2, a cBAF-specific component (Braun et al., 2017; Bergwell et al., 2024) (Fig. 6A and Supplemental Fig. 7B and C). The recruitment of Dpf2 with rapamycin to intron 3 of AR slightly increased AR-V7 mRNA level in 22Rv1 (Fig. 6B) and LNCaP (Supplemental Fig. 8E) cells. We hypothesized that the minimal increase in AR-V7 levels with Dpf2 recruitment in 22Rv1 cells was because this region is already maximally accessible. Therefore, we simultaneously treated cells with BD98 to inhibit cBAF association with chromatin and recruited Dpf2 to the locus (Supplemental Fig. 8F). Under these conditions, cBAF recruitment partially restored the BD98-induced reduction of AR-V7 (Fig. 6C) and RPB1-T4P (Fig. 6D). To confirm that these changes are dependent on the chromatin remodeling function of cBAF, we co-treated cells with BRM014 and rapamycin, which abrogated any effects of Dpf2 recruitment (Fig. 6C and D). These results suggest that the chromatin remodeling activity of cBAF at the transcriptional terminal region is required for AR-V7 expression.

**Figure 6.**
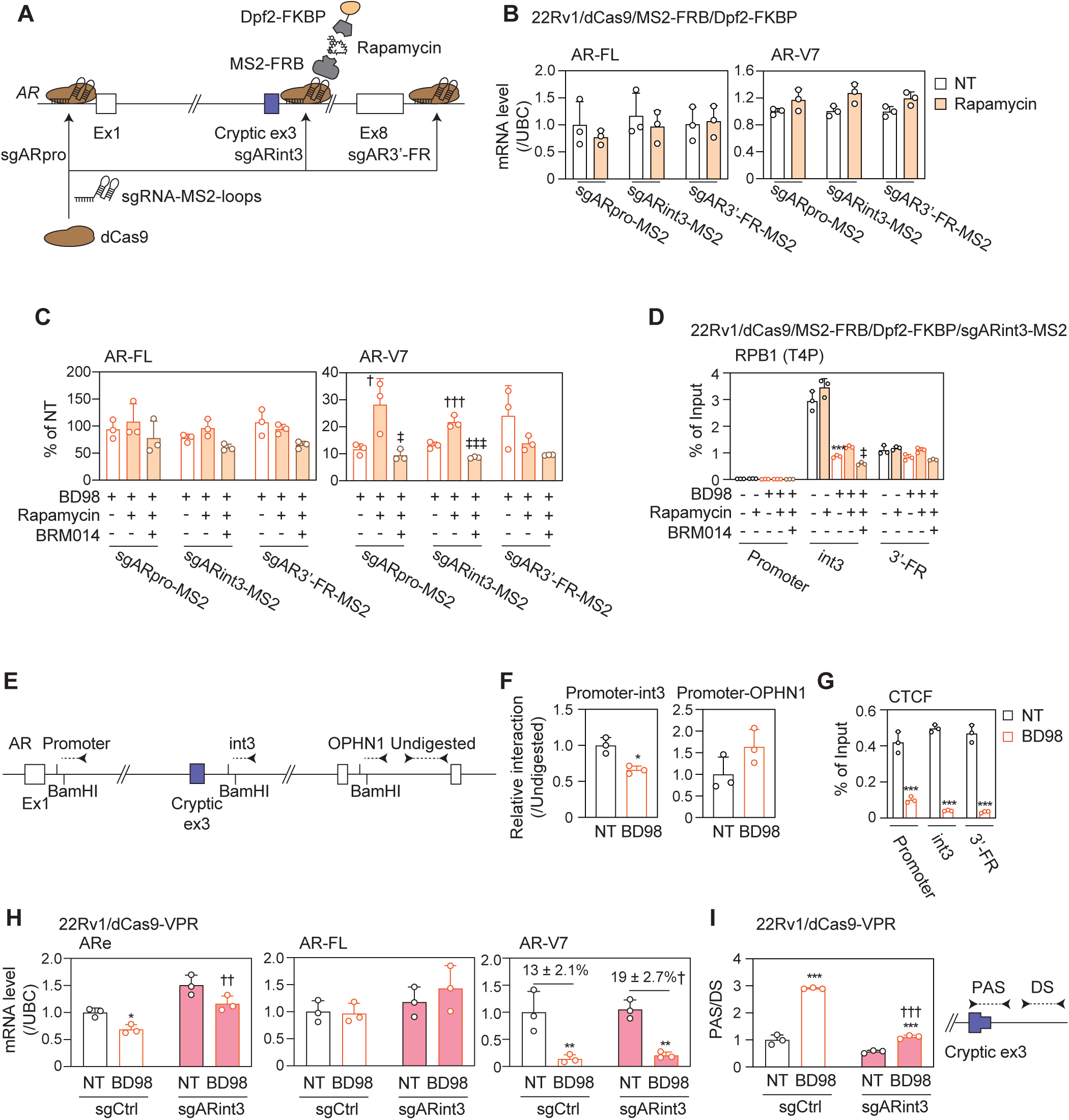
Effects of chromatin state on the transcriptional termination of AR-V7. (A) Dpf2 recruitment system. AR-FL and AR-V7 mRNA levels (B, C) and ChIP-qPCR for RPB1 (T4P) (D) of 22Rv1/dCas9/MS2-FRB/Dpf2-FKBP/sgRNA cells treated with 5 μM BD98, 5 nM rapamycin, and/or 5 μM BRM014 in FBS-containing medium for 96 hr. Each column represents the mean ± SD of three independent experiments. \*\*\**P* < 0.001, compared with NT, †*P* < 0.05 and †††*P* < 0.001, compared with BD98, ‡*P* < 0.05 and ‡‡‡*P* < 0.001, compared with BD98 and rapamycin. (E) Primers designed for the 3C assay. 3C assay (F) and ChIP-qPCR of CTCF (G) for 22Rv1 cells treated with 5 μM BD98 in FBS-containing medium for 48 hr. ARe, AR-FL, and AR-V7 levels (H) and the cleavage efficiency of the 3’ end of AR-V7 pre-mRNA (I) for 22Rv1/dCas9-VPR cells treated with 5 μM BD98 in FBS-containing medium for 96 hr. \**P* < 0.05, \*\**P* < 0.01, and \*\*\**P* < 0.001, compared with NT, †*P* < 0.05, ††*P* < 0.01, and †††*P* < 0.001, compared with sgCtrl

Unexpectedly, the recruitment of Dpf2 to the promoter region of *AR* also increased *AR-V7*, but not *AR-FL*, mRNA level (Fig. 6C). Public Hi-C data indicates that the AR promoter and the AR-V7 termination region are physically associated in 22Rv1 cells (Supplemental Fig. 9E); therefore, the recruitment of Dpf2 to the promoter region may also increase chromatin accessibility at intron 3. In LNCaP cells, in contrast, the promoter of AR does not interact with the AR-V7 termination region but instead interacts with a site in the *oligophrenin-1* (*OPHN1*) gene body (Supplemental Fig. 9G). We confirmed the interaction between the promoter and AR-V7 termination region by 3C assay in 22Rv1 cells (Fig. 6E, Supplemental Fig. 9H). Upon treatment with BD98, this interaction decreases and the interaction between the promoter and *OPHN1* increases (Fig. 6F). In addition, the binding of CTCF, a mediator of loop formation, to these regions was repressed by BD98 treatment (Fig. 6G). These results suggest that cBAF maintains chromatin loop formation at the *AR* gene.

From the POINT-seq we observed that BD98 treatment decreased signal of an antisense transcript near the transcriptional termination region of AR-V7 (Fig. 5A). This region exhibits enhancer-like characteristics, including a high ATAC signal, elevated H3K27ac levels, and cBAF binding. Because enhancer RNAs can be transcribed from intragenic enhancer regions to affect host gene expression (Cinghu et al., 2017), we examined whether this intragenic enhancer RNA (which we called ARe) affects AR-V7 expression using anti-sense oligonucleotides (ASO). Knockdown of ARe with ASO decreased AR-V7 mRNA and protein (Supplemental Fig. 8I and J) but not the levels of T4P bound to the region (Supplemental Fig. 8K). Treatment with BD98 decreased ARe levels, which were restored by transcriptional activation of intron 3 using a CRISPRa system to recruit dCas9-VPR. Restoration of AR-V7 mRNA levels, however, was limited (Fig. 6H), and T4P level was not changed by transcriptional activation of ARe (Supplemental Fig. 8L). Therefore, we evaluated 3’ processing of AR-V7 pre-mRNA by assessing the cleavage of the 3’-end of chromatin associated AR-V7 RNA. Using qRT-PCR, we compared the ratio of the mRNA around the alternate polyadenylation site (PAS) to the region downstream of cryptic exon 3. The cleavage of AR-V7 3’-end was impaired by BD98 treatment and restored by CRISPR activation (Fig. 6I), suggesting that the re-expression of ARe can rescue 3’ processing, but is not enough to rescue transcriptional termination. Collectively, cBAF remodels the chromatin environment to promote AR-V7 mRNA production by regulating transcriptional termination and 3’ processing.

### Global effects of BD98 on transcriptional termination

Next, we examined whether cBAF can facilitate transcriptional termination at additional genes by assessing the global impact of BD98 on transcriptional readthrough. From the POINT-seq data, upon BD98 treatment, 12% of genes (class II genes) had extended transcription downstream of the PAS whereas 88% (class I genes) did not (Fig. 7A and B). For class II genes, the RPB1 T4P signal around PAS was reduced and shifted slightly to the downstream side (Fig. 7C), suggesting that cBAF inhibition attenuates transcriptional termination. The ARID1B (Fig. 7D), SMARCA4 (Fig. 7E), and ATAC peaks (Fig. 7F) were mainly detected at the transcriptional start site (TSS) both in class I and II genes. Among the class II genes, approximately 200 have ARID1B peaks, with the majority in the gene body (Fig. 7G), for example, *FNBP1L* (Fig. 7H). BD98 treatment decreased ARID1B and ATAC peaks and shifted the POINT and T4P signals downstream. These results indicate that cBAF-mediated remodeling around PAS can function as a gatekeeper of RNA polymerase II and affect transcriptional termination, primarily in an indirect manner through enhancer activation and chromatin looping.

**Figure 7.**
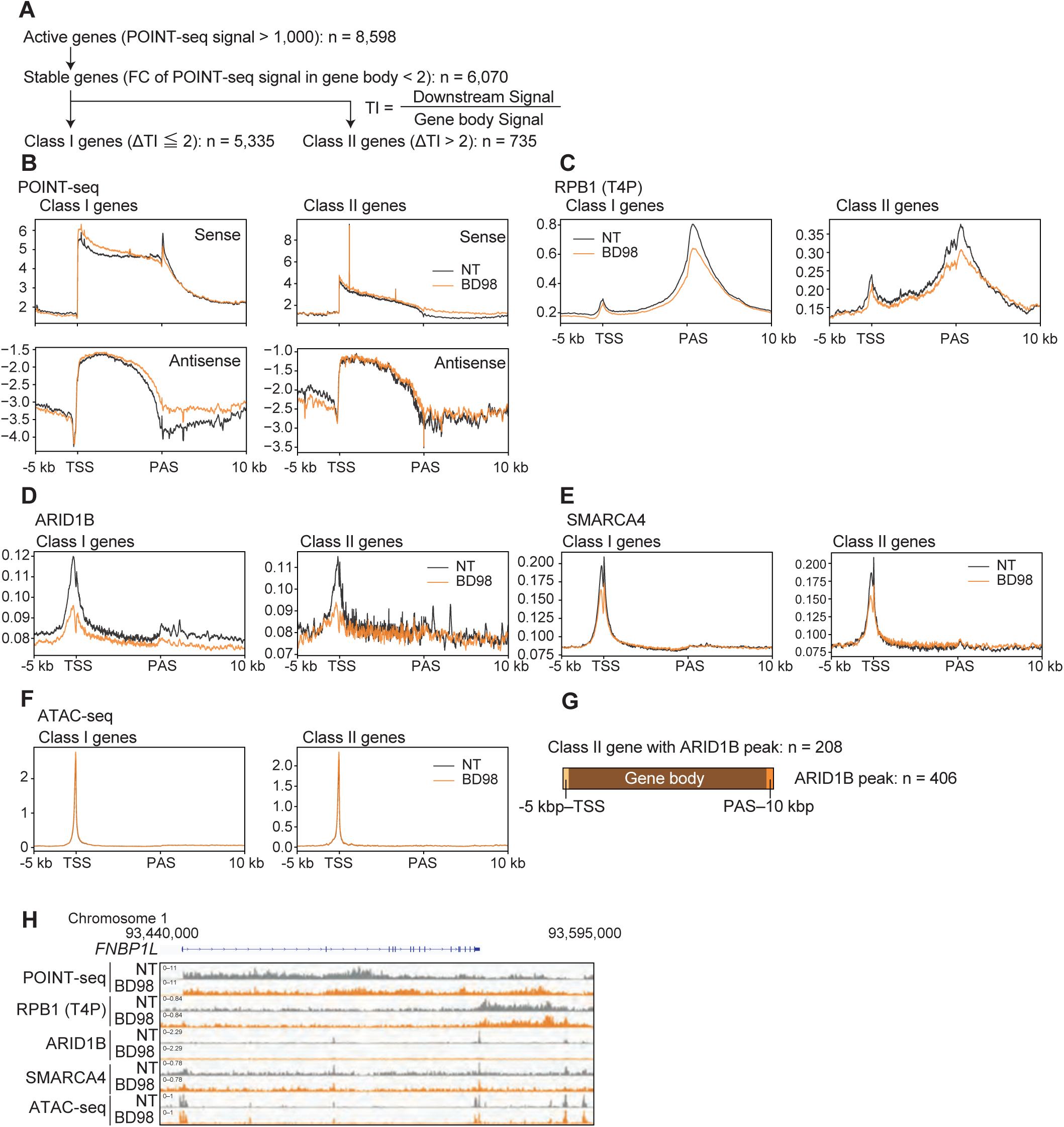
Effects of BD98 treatment on the global transcriptional termination of AR-V7. (A) 22Rv1 cells were treated with 5 μM BD98 in FBS-containing medium for 48 hr. Active genes were categorized into class I and II genes based on TI. Metagene plots (mean of replicates) of POINT-seq (B), ChIP-seq for RPB1 (T4P) (C), ARID1B (D), and SMARCA4 (E), and ATAC-seq (F) are shown for Class I and II genes. (G) Distribution of ARID1B peaks in class II genes. (H) Genomic tracks of POINT-seq and ChIP-seq (RPB1 (T4P), ARID1B, and SMARCA4) at *FNBP1L*. Each area represents the mean of two (ChIP-seq and POINT-seq) or three (ATAC-seq) independent experiments.

## Discussion

Here we demonstrate that cBAF chromatin remodeling complex is a regulator of AR-V7 levels in CRPC cells, a role that is not shared with other SWI/SNF subcomplexes. Previous studies established that the production of AR-V7 mature mRNA is regulated by post-transcriptional splicing (Liu et al., 2014; Fan et al., 2018; Kawamura et al., 2019; Brittain et al., 2026). In addition, the CPSF1, polyadenylation factor, has been shown to regulate AR-V7 expression (Van Etten et al., 2017). The polyadenylation signal within cryptic exon 3 is present prior to CRPC development, suggesting that AR-V7 is determined by its utilization rather than its acquisition. Our data indicate that cBAF maintains chromatin accessibility at this site, contributing to RNA polymerase II pausing downstream of cryptic exon 3 (Fig. 5A). Perhaps related, a recent report demonstrated that knockdown of cyclin-dependent kinase 12 (CDK12), a key regulator of transcriptional elongation (Tellier et al., 2020), also increases AR-V7 expression in prostate cancer cells (Sun et al., 2022). Given that CDK12 suppresses intronic polyadenylation (Dubbury et al., 2018), accelerated transcriptional elongation may enable RNA polymerase II to read through the cryptic exon 3 region, thereby suppressing premature termination events associated with AR-V7 generation. Thus, suppression of AR-V7 expression in normal prostate cells requires coordination by chromatin structure, transcriptional dynamics, and the RNA processing machinery. During the progression from normal prostate to CRPC, there is an increase in chromatin relaxation (Braadland PR and Urbanucci A, 2019), enhancer activation (Shuai Y and Huang, 2025), and chromatin looping (Ramanand et al., 2020), which facilitates transcriptional plasticity and therapeutic resistance. While we have shown that recruiting cBAF to this locus in hormone naïve LNCaP cells can increase AR-V7 levels slightly (Supplemental Fig. 8E), additional alterations in chromatin structure are likely required for cBAF to promote chromatin accessibility and increased AR-V7 expression during the development of castration resistance.

Transcription termination in eukaryotes has long been considered primarily controlled by cis-acting sequence elements, particularly the polyadenylation signal at the 3’ end of genes (Proudfoot, 2019). In addition to cis-acting RNA processing signals, histone modifications have been implicated in regulating transcription termination efficiency. For example, histone H3 lysine 36 trimethylation (H3K36me3) covers the entire body of active genes, peaking at the PAS (Edmunds et al., 2008), and knockout of SET domain containing 2 (SETD2) causes transcriptional readthrough of a subset of genes (Kopczyńska et al., 2026). While much remains to be uncovered regarding epigenetic control of transcriptional termination, the present study provides new insight into region-dependent regulation of transcriptional termination site usage through enhancer activation.

The SWI/SNF complex is a key transcriptional regulator in many cell types, modulating chromatin accessibility at promoters and enhancers to facilitate transcription factor binding and RNA polymerase II recruitment (Ryan et al., 1998). Additional studies have expanded this framework by implicating SWI/SNF complexes in transcriptional elongation and gene body regulation (Schwabish and Struhl, 2007; Cornejo et al., 2024; Morse et al., 2024). Our results further extend the functional scope of gene regulation by SWI/SNF to include the regulation of transcriptional termination by facilitating intronic enhancers that regulate transcriptional kinetics. In the case of AR-V7, cBAF regulates transcriptional termination and 3’ processing through chromatin remodeling, chromatin loop formation, and transcriptional activation of intragenic enhancer RNA (Fig. 6). Our POINT-Seq data indicates that this mechanism may be broadly applicable to other genes in 22Rv1 cells (Fig. 7); however, further studies will be required to validate this role for cBAF at additional genes and in other cell types.

Our findings provide important translational implications for the treatment of CRPC. Although AR-V7 is a well-established driver of resistance to anti-androgen therapies, its truncated structure and lack of a ligand-binding domain have made it difficult to directly target pharmacologically (Wang et al., 2017), although efforts are underway to target N-terminal and DNA-binding regions of AR for novel proteolysis-targeting chimeras that induce AR-V7 degradation (Ma et al., 2022; Ha et al., 2025). As an alternative approach, SWI/SNF is a well-established target for inhibiting AR-mediated transcription in CRPC (Ordonez-Rubiano et al, 2024). SMARCA2/4 degraders are effective against CRPC and can even resensitize VCAP xenografts to AR antagonists (Xiao et al., 2022). This study demonstrates cBAF inhibition exerts this anti-proliferative effect on CRPC cells (Fig. 3D and Supplemental Fig. 1F) not only by inhibiting AR-FL binding to its target region (Fig. 3E) but also by suppressing *AR-V7* expression (Fig. 2E and F). Together, these findings position cBAF inhibition as a therapeutic strategy capable of overcoming AR-V7-mediated resistance in CRPC.

## Limitations of the Study

Our conclusions are based primarily on *in vitro* models of prostate cancer, principally the 22Rv1 cell line, with additional validation in VCaP and LNCaP cells. The pharmacological effects of cBAF inhibition on AR-V7 and CRPC *in vivo* remain to be established. Future studies employing relevant *in vivo* models, including patient-derived xenografts and immunocompetent mouse models, will be critical to validate the therapeutic potential of targeting cBAF in AR-V7-positive CRPC. Second, our genome-wide analysis identified a subset of genes (class II) that undergo cBAF-dependent transcriptional readthrough upon BD98 treatment, but it remains unknown whether these loci have similar chromatin features to the transcriptional termination region of AR-V7, such as enhancer-like activity and looping formation. It will be necessary to determine what chromatin state features define cBAF-regulated termination regions genome-wide.

## Abbreviation

AR: Androgen receptor
AR-FL: Full length androgen receptor
ARID: AT-rich interactive domain-containing protein
AR-V7: Androgen receptor splice variant 7
ASO: Antisense oligonucleotide
ATAC: Assay for transposase-accessible chromatin
BAF: BRG1/BRM-associated factor
BRD: Bromodomain containing
BRG1: Brahma-related gene 1
BRM: Brahma
BSD: blasticidin S deaminase
BT: BromoTag
3C: Chromosome conformation capture
cBAF: Canonical BAF
CBX2: Chromobox 2
ChIP: Chromatin immunoprecipitation
CE: Cytoplasmic extract
CPSF: Cleavage and polyadenylation specificity factor
CRISPR: Clustered regularly interspaced short palindromic repeat
CRISPRa: CRISPR activation
CRPC: Castrate resistant prostate cancer
CTCF: CCCTC-binding factor
CUT&RUN: Cleavage under targets & release using nuclease
Cytoplasmic: extract
CSS: Charcoal-dextran stripped FBS
CTD: *C*-terminal domain
DEG: Differentially expressed genes
DHT: Dihydrotestosterone
DMEM: Dulbecco’s Modified Eagle Medium
Dox: Doxycycline
Dpf2: Double PHD fingers 2
EDTA: Ethylenediaminetetraacetic acid
EGTA: Ethylene glycol tetraacetic acid
EMT: Epithelial mesenchymal transition
ENZA: Enzalutamide
ex: Exon
F12: Nutrient mixture F-12
FAIRE: Formaldehyde-assisted isolation of regulatory elements
FBS: Fetal bovine serum
FC: Fold change
FKBP: FK506-binding protein
FR: Flanking region
FRB: FKBP-rapamycin binding domain of mTOR
FOXA1: forkhead box A1
GBAF: GLTSCR1/1L-BAF
GLTSCR: Glioma tumor suppressor candidate region
HA: Hemagglutinin
HEPES: 4-(2-Hydroxyethyl)-1-piperazineethanesulfonic acid
H3K36me3: histone H3 lysine 36 trimethylation
HNRNPF: Heterogeneous nuclear ribonucleoprotein F
int: Intron
JMJD1A: Jumonji domain containing 1A
KLK: kallikrein related peptidase
MEM: NEAA Minimum essential medium non-essential amino acids
mTORC1: Mechanistic/mammalian target of rapamycin complex 1
NE: Nuclear extract
NP-40: Nonidet P-40
NT: Non-treatment
OPHN1: Oligophrenin-1
RPMI: Roswell Park Memorial Institute
PAS: Polyadenylation site
PBAF: Polybromo-associated BAF
PBRM1: Polybromo-1
PBS: Phosphate-buffered saline
PC: prostate cancer
PCR: Polymerase chain reaction
PNUTS: Phosphatase 1 nuclear targeting subunit
PP: Protein phosphatase
Pre-mRNA: Precursor mRNA
qPCR: Quantitative polymerase chain reaction
RPB1: RNA polymerase II subunit B1
RT: Reverse transcription
RPM: reads per million
SAHA: Suberoylanilide hydroxamic acid
SDS: Sodium dodecyl sulfate
sgRNA: Single guide RNA
shRNA: Short hairpin RNA
TBP: TATA-binding protein
TI: Termination index
SMARC: SWI/SNF related BAF chromatin remodeling complex subunit
SWI/SNF: Switch/sucrose non-fermentable
TE: Tris-EDTA
TSS: Transcriptional start site
VHL: von Hippel-Lindau
VPR: VP64-p65-Rta

## Acknowledgements

The authors thank Dr. Hiroshi Kimura for providing and Dr. Diana Hargreaves for providing GIPZ Lentiviral shRNAs and the RNA polymerase II antibody (MABI601), respectively. The authors appreciate Dr. Takayuki Nojima for valuable advice and guidance about POINT-seq. The authors thank the Purdue Genomics Core for technical support for sequencing analysis.

## Availability of Data

Sequencing data generated in this study have been deposited in the NCBI Gene Expression Omnibus (GEO) under the following accession numbers: RNA-seq (GSE350177, GSE350178, GSE350179, and GSE350180), ChIP-seq (GSE350176), CUT&RUN (GSE350169), ATAC-seq (GSE350168), POINT-seq (GSE350170). The mass spectrometry data have been deposited to the ProteomeXchange Consortium via the jPOST repository (Okuda et al., 2025) under accession numbers PXD085590 (ProteomeXchange) and JPST004992 (jPOST).

## Author contributions

Conceptualization, Data curation, Formal analysis, Investigation, Project administration, Validation, Visualization, and Writing – original draft, M.N.; Investigation, S.U.; Resources, J.G.K.; Investigation, W.A.T.; Conceptualization, Data curation, Funding acquisition, Project administration, Supervision, and Writing – review & editing, E.C.D.

---

**Supplemental Figure 1.**
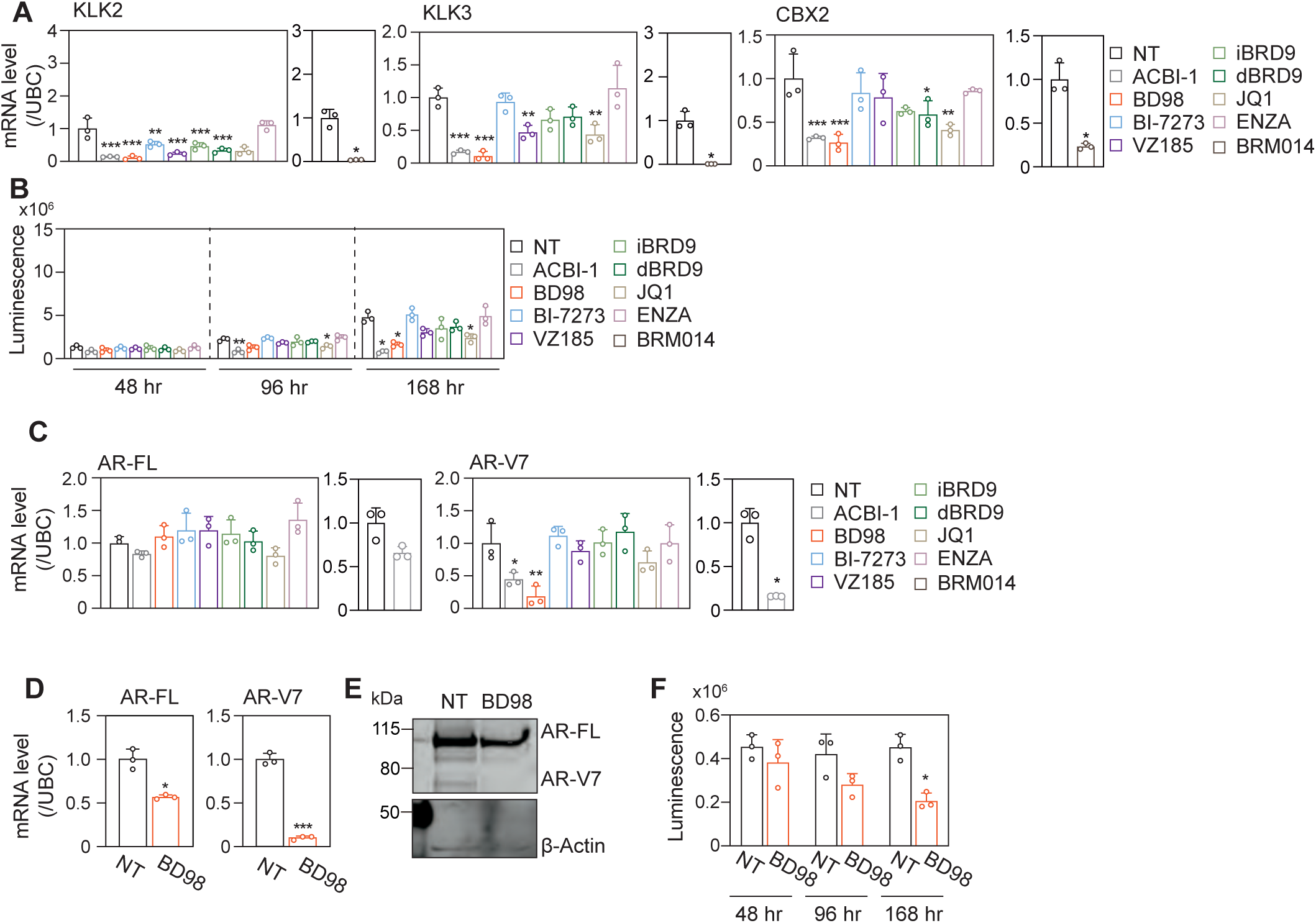
Effects of SWI/SNF inhibitors on AR-FL- and AR-V7-mediated transcriptional regulation and their expression. KLK2, KLK3, CBX2 (A), AR-FL, and AR-V7 (C) mRNA levels, viability (B and F), and AR protein level (E) of 22Rv1 (A–C) and VCaP cells (D–F) treated with 0.5 μM ACBI-1, 5 μM BD98, 5 μM BI-7273, 0.5 μM VZ-185, 5 μM iBRD9, 0.5 μM dBRD9, 5 μM JQ1, 10 μM ENZA, or 5 μM BRM014 in CSS-containing medium for 48 (B and F), 96 (A–F), or 168 hr (B and F). Representative Western blots of three independent experiments are shown. Each column represents the mean ± SD of three independent experiments. \**P* < 0.05, \*\**P* < 0.01, and \*\*\**P* < 0.001, compared with NT.

**Supplemental Figure 2.**
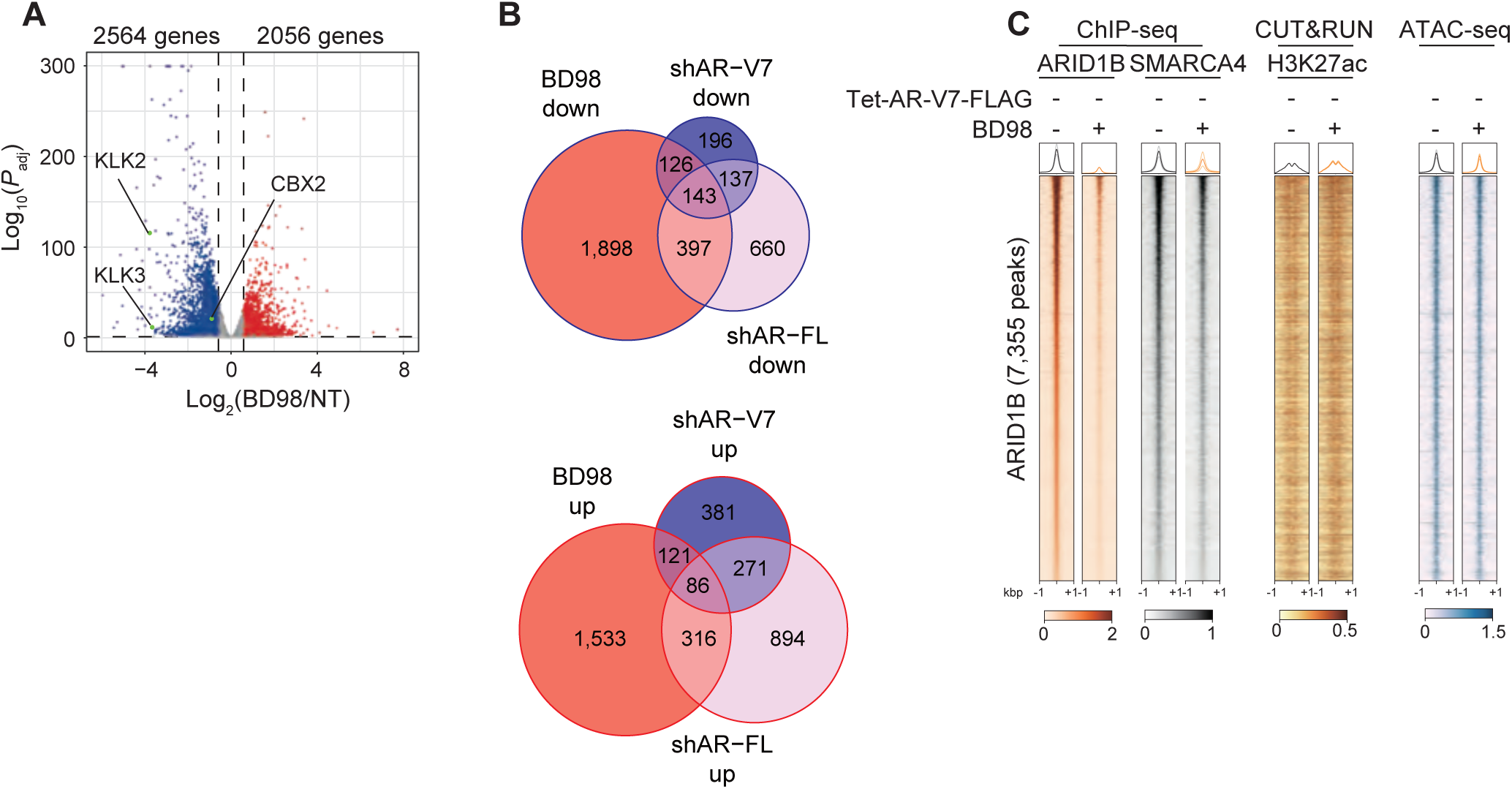
Treatment with BD98. (A) RNA-seq for 22Rv1/Tet-Empty cells treated with 5 μM BD98 in CSS-containing medium for 96 hr (three independent experiments). In the volcano plot, genes whose expression was significantly decreased or increased (*P*_adj_ < 0.05 and FC > 1.5) by BD98 treatment are shown in blue or red, respectively. (B) Overlap of DEGs with treatment BD98 and those with transduction sh*AR-FL* or sh*AR-V7* knockdown (Fig. 1C). (C) ChIP-seq for ARID1B and SMARCA4 (two independent experiments), CUT&RUN for H3K27ac (two independent experiments), and ATAC-seq (three independent experiments) for 22Rv1 cells treated with 5 μM BD98 in FBS-containing medium for 48 hr. Metagene plots (faint line, each replicate; dark line, mean of replicates) and heatmaps (mean of replicates) are shown for ARID1B-bound regions. (D) Motif analysis for AR-FL-specific, AR-V7-specific, and their common regions.

**Supplemental Figure 3.**
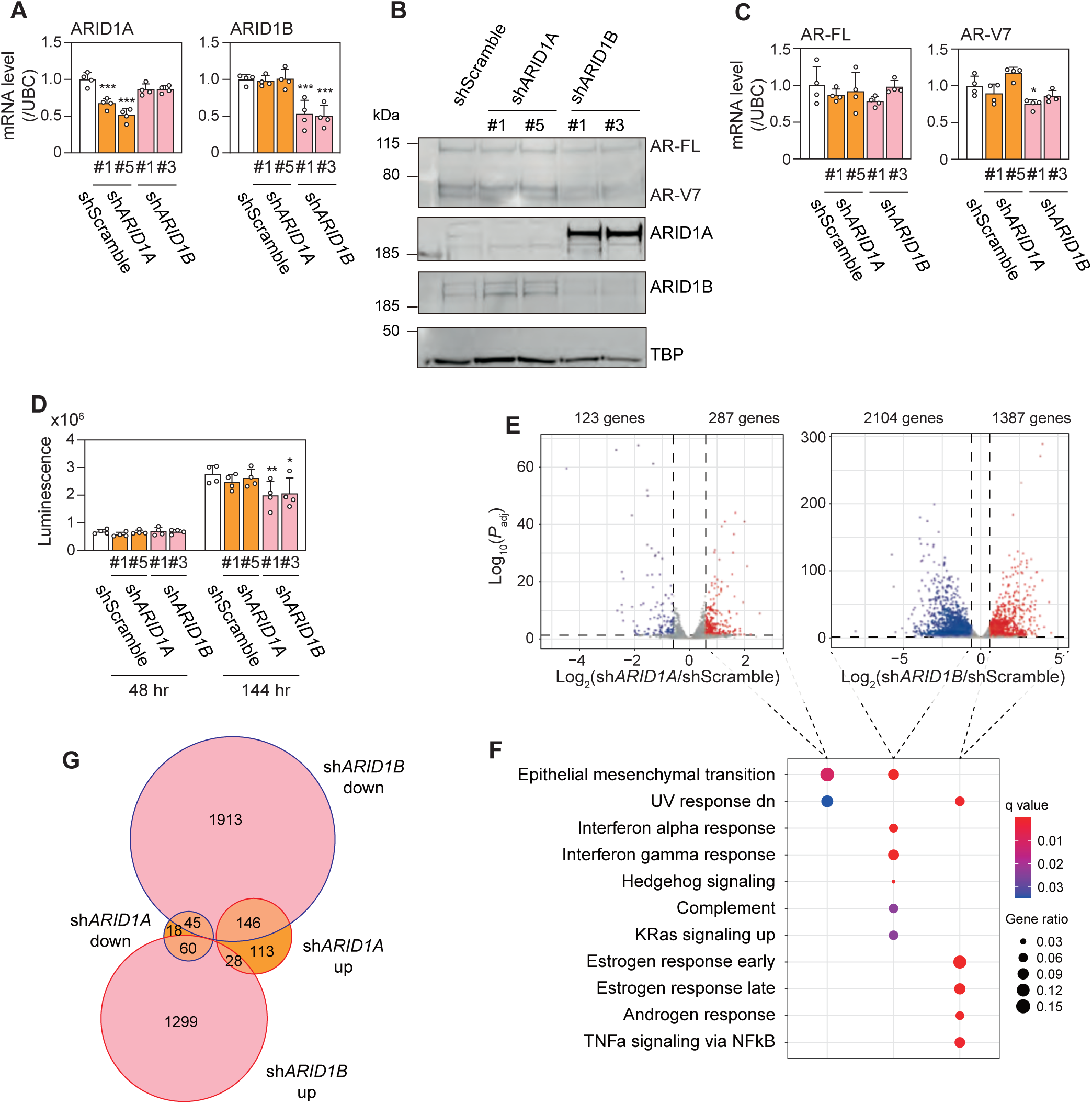
Knockdown of ARID1A or ARID1B by shRNA. ARID1A, ARID1B (A), AR-FL, and AR-V7 (C) mRNA, protein levels (B), viability (D), and RNA-seq for 22Rv1 cells transduced with sh*ARID1A* or sh*ARID1B* and incubated in CSS-containing medium for 72 hr. Representative Western blots of three independent experiments are shown. Each column represents the mean ± SD of three or four independent experiments. \**P* < 0.05, \*\**P* < 0.01, and \*\*\**P* < 0.001, compared with shScramble. (E) In the volcano plot, genes whose expression was significantly decreased or increased (*P*_adj_ < 0.05 and FC > 1.5) by transduction with sh*ARID1A* or sh*ARID1B* are shown in blue or red, respectively. (F) The significantly enriched MSigDB Hallmark gene sets in DEGs are shown. (G) Overlap of the DEGs transduction with ARID1A, ARID1B knockdown and those with BD98 treatment (Supplemental Fig. 2A).

**Supplemental Figure 4.**
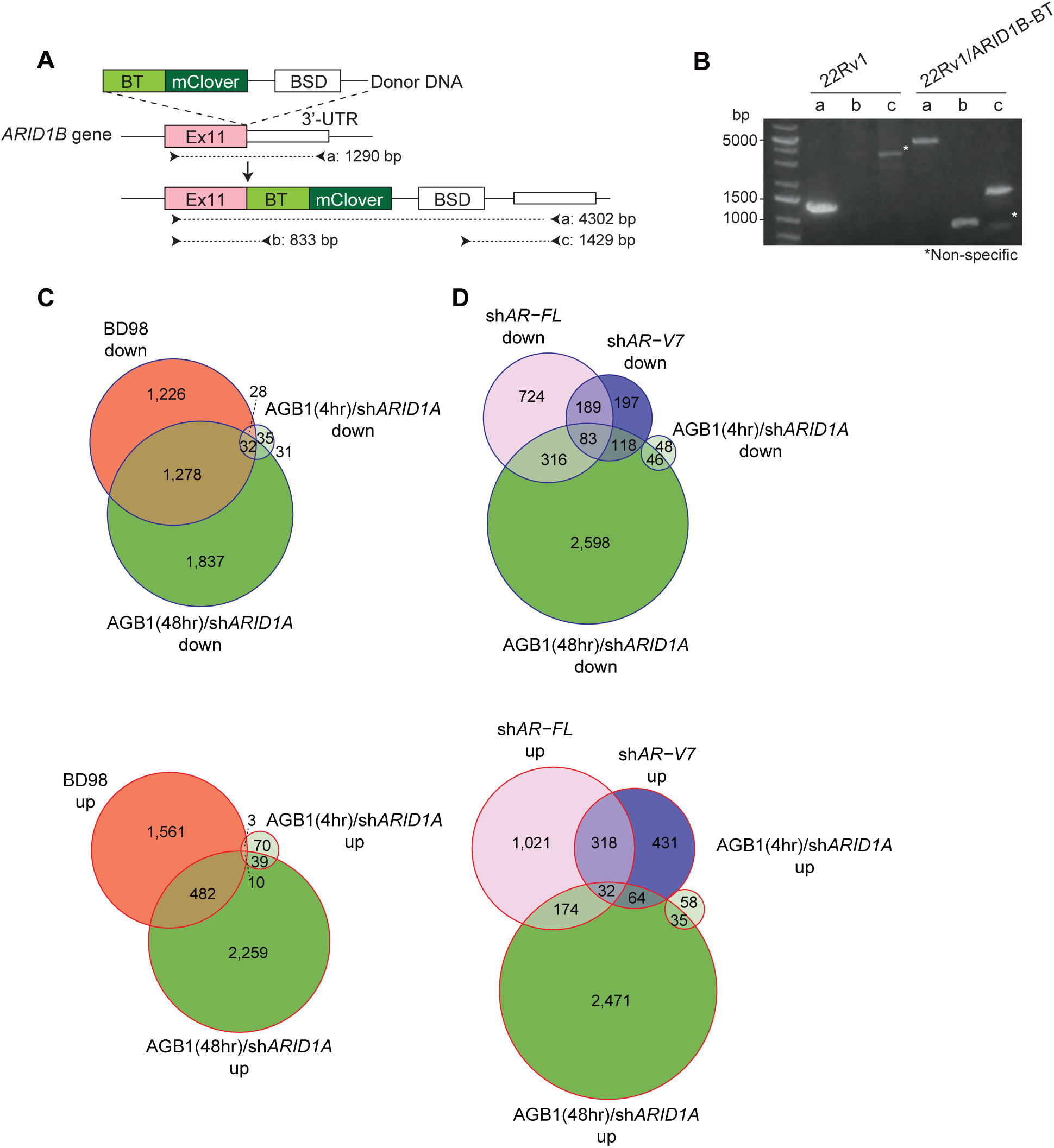
ARID1B degron system. (A) Knock-in of BT into the *ARID1B* gene. (B) Genotyping of 22Rv1/1B-BT cells. (C) Overlap of the DEGs with knockdown of ARID1A and/or ARID1B by treatment with AGB1 for 4 hr (Down, Clusters 1–4; Up, Clusters 5–8 in Fig. 4D) or 48 hr (Down, Clusters 1, 2; Up, Clusters 3, 4 in Fig. 4F) and those with BD98 treatment (Supplemental Fig. 2A) (C) or those with transduction with sh*AR-FL* or sh*AR-V7* (Fig. 1C) (D).

**Supplemental Figure 5.**
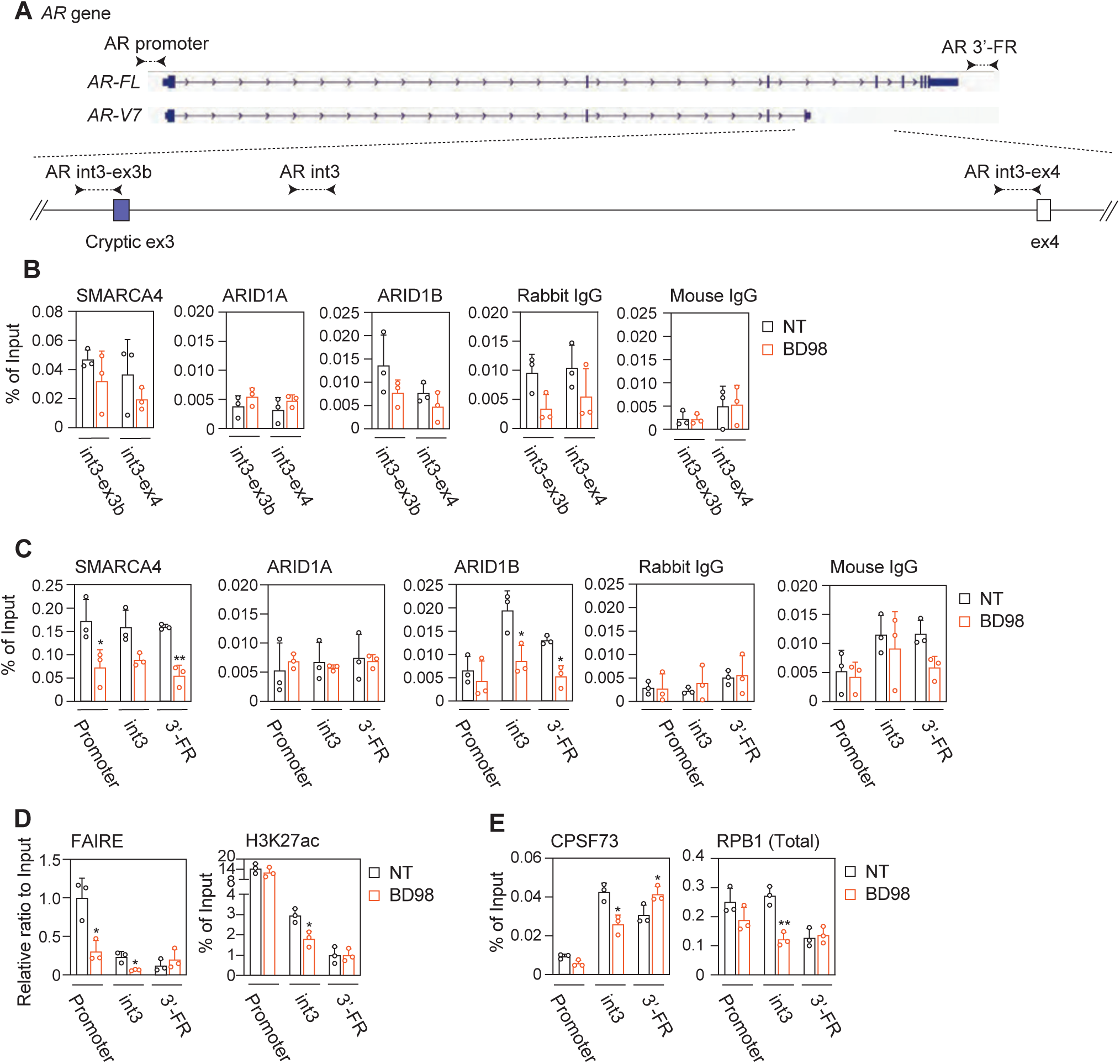
Effects of BD98 treatment on transcriptional termination of AR-V7. (A) Structure of the *AR* gene and locations of primers designed for ChIP-qPCR. ChIP-qPCR of SMARCA4, ARID1A, ARID1B (B, C), H3K27ac (D), CPSF73, and RPB1 (Total) (E), and FAIRE assay (D). Each column represents the mean ± SD of three independent experiments. \**P* < 0.05 and \*\**P* < 0.01, compared with NT. ex, exon; int, intron.

**Supplemental Figure 6.**
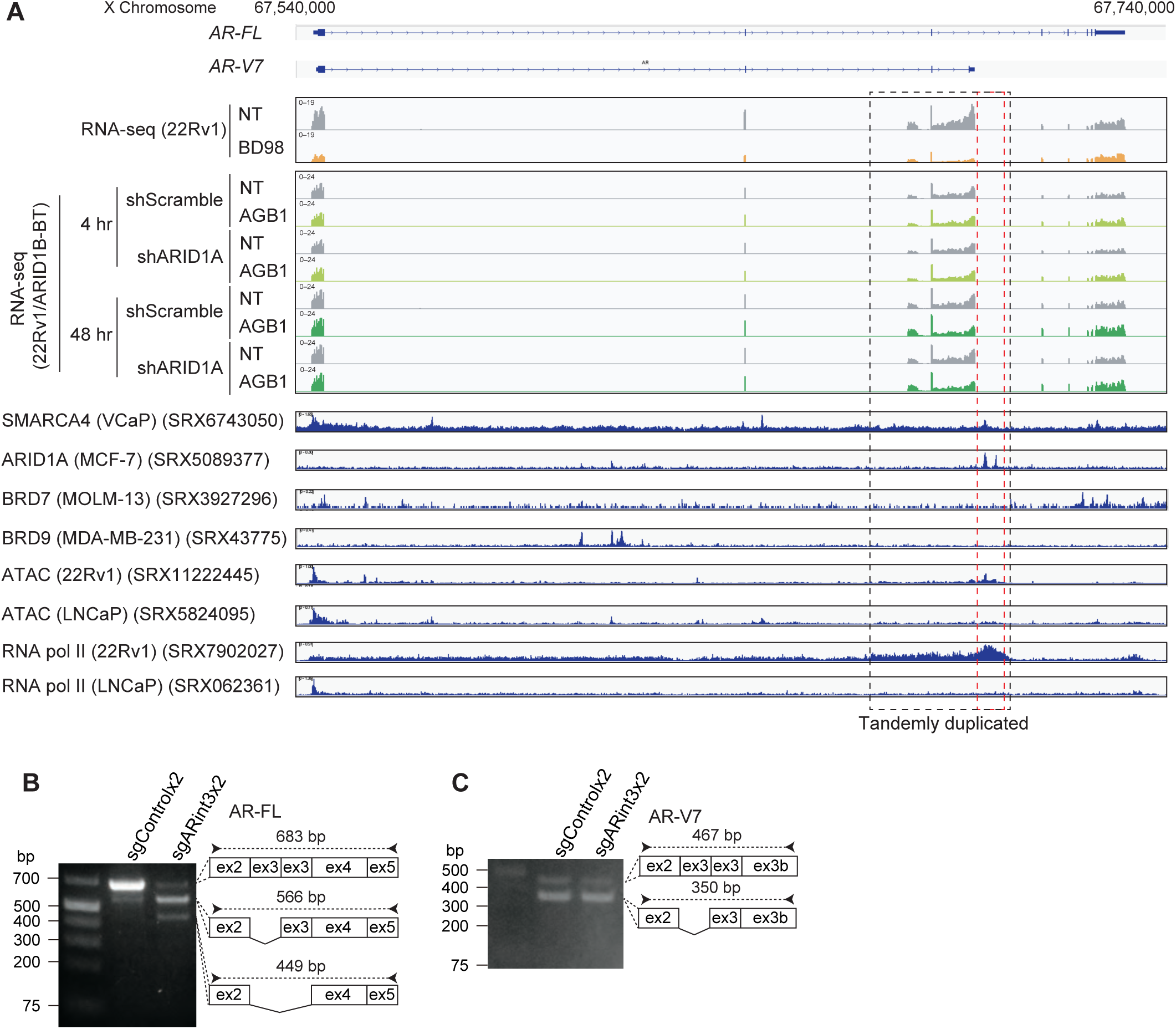
Transcriptional termination of AR-V7. (A) Genomic tracks of RNA-seq at the *AR* gene in 22Rv1 cells treated with 5 μM BD98 in CSS-containing medium for 96 hr and 22Rv1/ARID1B-BT cells transduced with shARID1A and treated with 200 nM AGB1 for 4 or 48 hr. Public ChIP-seq data obtained via ChIP-Atlas (Zou et al., 2022). Splicing patterns of AR-FL (B) and AR-V7 (C) in 22Rv1 cells transfected with pX333/sgARint3×2 in FBS-containing medium for 72 hr.

**Supplemental Figure 7.**
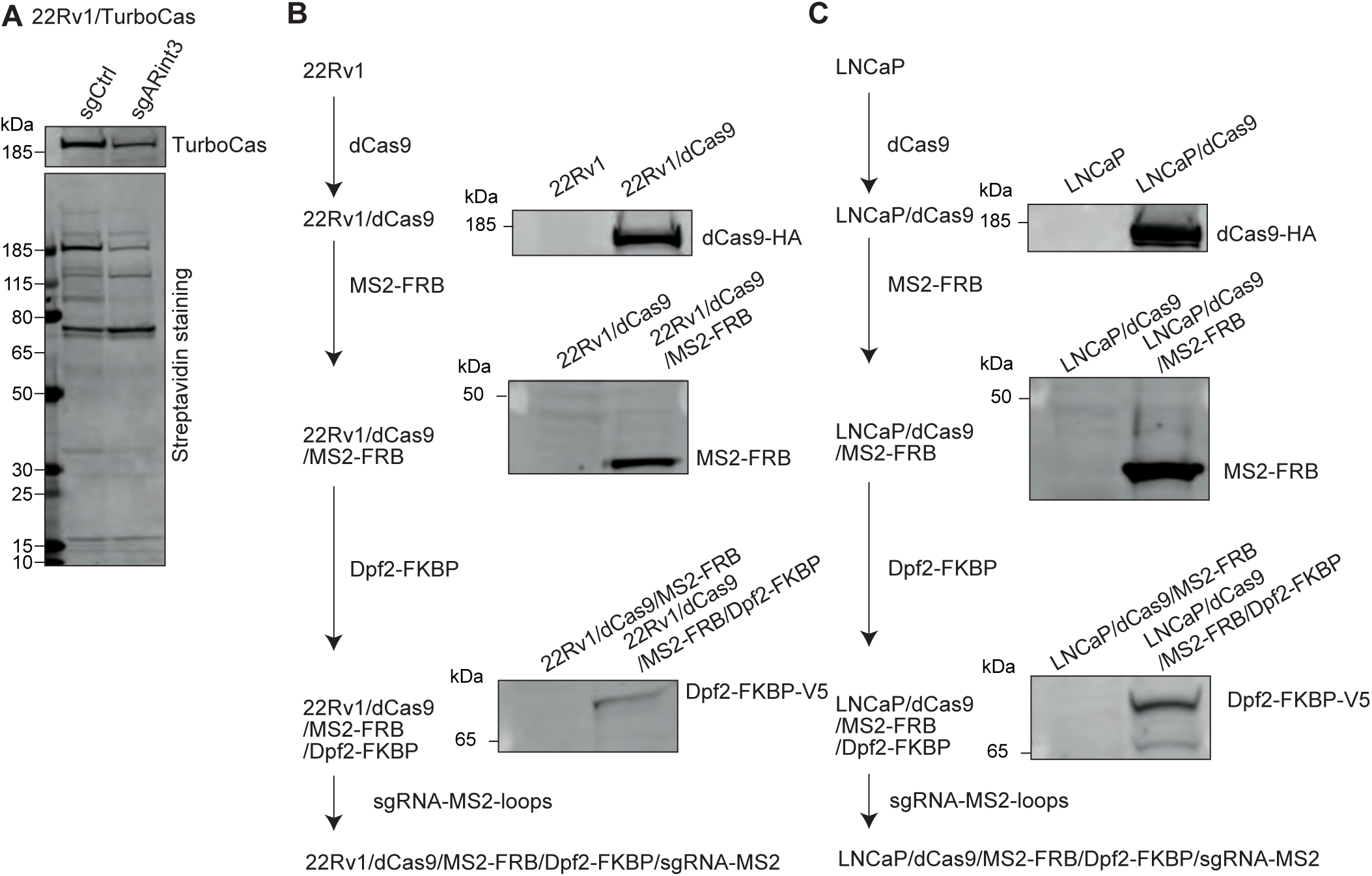
Establishment of the CRISPR recruitment system. (A) Western blot for TurboCas and streptavidin staining of 22Rv1/TurboCas. 22Rv1 (B) and LNCaP (C) cells were sequentially transduced with dCas9, MS2-FRB, Dpf2-FKBP, and sgRNA-MS2-loops.

**Supplemental Figure 8.**
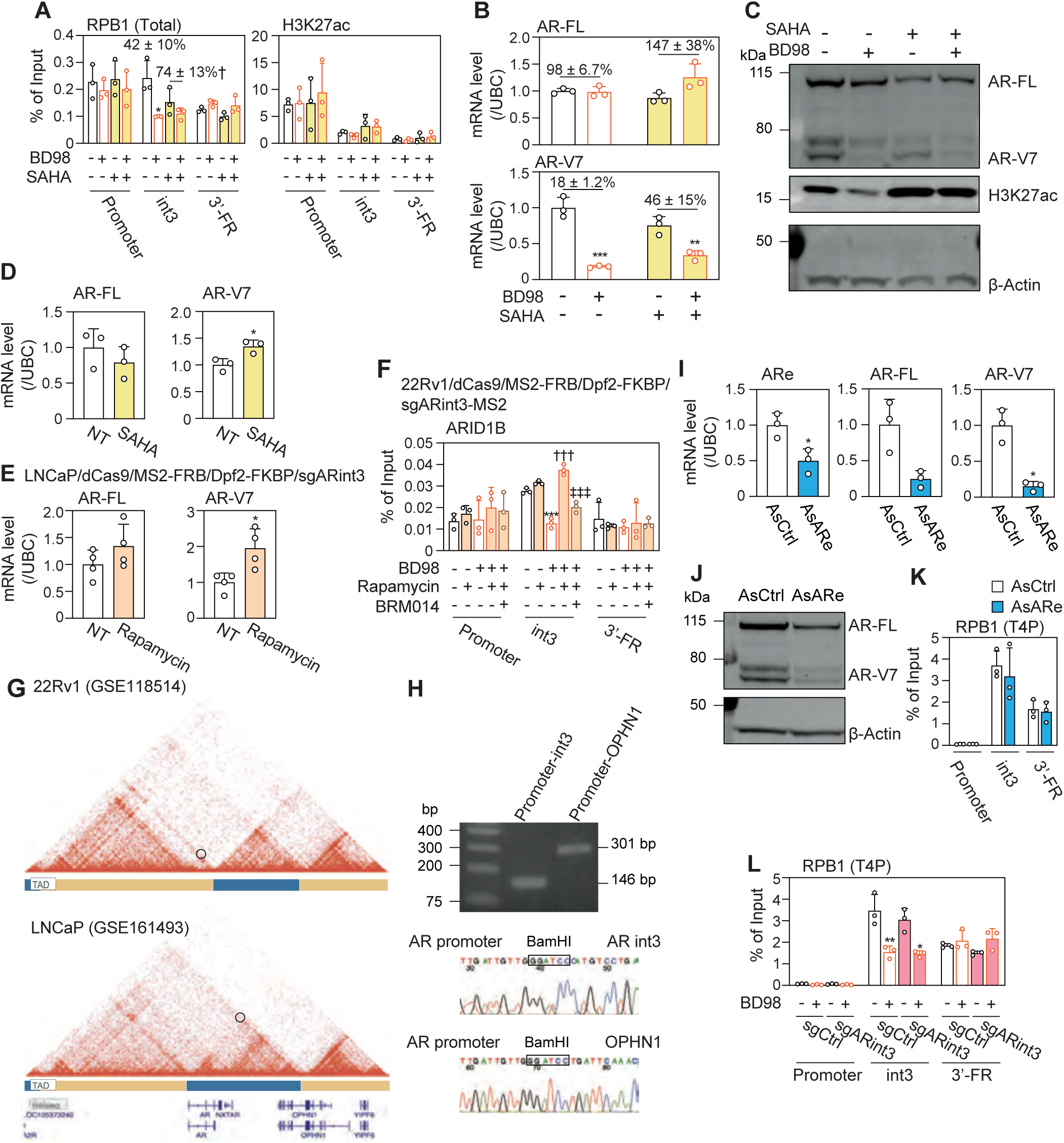
Effects of chromatin environment on AR-V7 expression. ChIP-qPCR of RPB1 and H3K27ac (A) and AR-FL, AR-V7, and H3K27ac mRNA (B) and protein (C) levels for 22Rv1 (A–C) and LNCaP (D) cells treated with 5 μM SAHA and/or 5 μM BD98 in FBS-containing medium for 96 hr. (E) AR-FL and AR-V7 mRNA levels in LNCaP/dCas9/MS2-FRB/Dpf2-FKBP/sgARint3 cells treated with 5 nM rapamycin in FBS-containing medium for 96 hr. Each column represents the mean ± SD of three or four independent experiments. \**P* < 0.05, \*\**P* < 0.01, and \*\*\**P* < 0.001, compared with NT, †*P* < 0.05, compared with SAHA(-), (F) ChIP-qPCR for ARID1B of 22Rv1/dCas9/MS2-FRB/Dpf2-FKBP/sgARint3-MS2 cells treated with 5 μM BD98, 5 nM rapamycin, and/or 5 μM BRM014 in FBS-containing medium for 96 hr. \**P* < 0.05, \*\**P* < 0.01, and \*\*\**P* < 0.001, compared with NT, †††*P* < 0.001, compared with BD98, ‡‡‡*P* < 0.001, compared with BD98 and rapamycin. (G) Public Hi-C data obtained via 3D Genome Browser 2.0 (Yu et al., 2026). (H) PCR products and Sanger sequencing of the 3C assay for 22Rv1 cells. ARe, AR-FL, and AR-V7 mRNA (I) and protein (J) levels, and ChIP-qPCR of RPB1(T4P) (K) for 22Rv1 cells transfected with 100 nM AsARe for 24 hr. \**P* < 0.05, compared with AsARe. (L) ChIP-qPCR of RPB1(T4P) for 22Rv1/dCas9-VPR cells treated with BD98 in FBS-containing medium for 96 hr. \**P* < 0.05 and \*\**P* < 0.01 with NT. Representative Western blots of three independent experiments are shown.

